# *NicheProt*: Cell-type-resolved proteomics of tissue compartments

**DOI:** 10.1101/2025.10.23.684231

**Authors:** Yi-Chien Wu, Dylan Schwartz, Elie Abi Khalil, Aditi Upadhye, Jalees Rehman, Steve Seung-Young Lee

## Abstract

Spatial proteomics uncovers the molecular underpinnings of cellular function in intact tissues. Laser capture microdissection coupled with mass spectrometry enables comprehensive proteomic profiling of selected tissue regions, but typically does not support cell-type-specific proteomic analysis. We present *NicheProt*, a 3D optical microscopy-guided, photobleaching-mediated cell barcoding approach for isolating intact specific cell types from defined microanatomical tissue compartments or niches. Using sequential bottom-up proteomic analysis, we defined two distinct phenotypes of CD11c⁺ dendritic cells based on their spatial locations in the inflamed mouse spleen. These two compartment-specific dendritic cell populations were characterized by proteomic signatures differing in the levels of 54 proteins. This 3D tissue microscopy-guided method offers cell-type and microregion-resolved proteomic analysis, facilitating the proteomic discovery of previously unrecognized cell subtypes and their functional roles in distinct tissue compartments.

## Main

The functional integrity of tissue depends on the precise spatial organization of diverse cell types and their dynamic interactions [1]. This spatial architecture governs tissue homeostasis, orchestrates immune responses, and shapes disease progression [2]. Biomolecules such as DNA, RNA, and proteins execute cellular functions, thus serving as the key readouts to decode complex cell biology and tissue organization [3].

In recent years, spatial “omics” approaches have emerged as transformative platforms, enhancing our understanding of developmental biology, pathogenesis, and therapeutic responses because they resolve functional and phenotypic differences between cells in specific tissue compartments. Among these, spatial transcriptomics enables RNA profiling in the tissue context and achieves subcellular resolution through advanced techniques, such as multiplexed *in situ* hybridization and sequencing platforms [4, 5]. A notable example is the commercialized Visium/Xenium platform by 10x Genomics, which captures mRNA transcripts *in situ* using a barcoded oligonucleotide array slide followed by downstream mRNA sequencing [6–8]. When combined with single-cell RNA sequencing, spatial transcriptomics further offers a detailed cellular landscape of tissue samples [9]. However, RNA levels do not always correlate with protein abundance due to translational regulation [10–12]. Since proteins are the primary effector molecules of cellular function, spatial proteomics offers a more direct window into cellular mechanisms in the tissue context.

Various spatial proteomics technologies have recently been developed, many of which rely on antibody-based strategies for targeted protein detection. These include metal-tagged antibodies (i.e., MIBI [13–15] and IMC [16]), oligonucleotide-conjugated antibodies (i.e., CODEX [17, 18], DBiT-seq [19], GeoMx DSP [20], and SUM-PAINT [21]), and fluorescent antibodies (i.e., cyclic-imaging-based proteomics [22–24]). While powerful, these methods are limited by their dependence on prior knowledge of protein targets and the restricted number of proteins that can be simultaneously analyzed. In contrast, mass spectrometry (MS) enables untargeted and comprehensive proteome profiling without antibody labeling. Among various MS-based spatial proteomics approaches, laser capture microdissection (LCM) coupled with MS analysis remains one of the most widely used techniques. This method is often used to isolate mesoscopic tissue regions composed of mixed cell types from thin formalin-fixed paraffin-embedded (FFPE) tissue sections (< 10 µm thick). In LCM, integration with immunofluorescence (IF) staining can help to localize and isolate specific cell types within a tissue section [25, 26]. However, this approach remains challenging, as it requires manual dissection of individual cells, which is labor-intensive and prone to contamination from adjacent unwanted cell types or extracellular matrix (ECM). These challenges are further exacerbated when isolating irregularly shaped (e.g., neurons and endothelial cells) or small cells located in densely packed areas (e.g., immune cells in lymphoid tissues). High-energy laser cutting in LCM can also damage the cell membrane, leading to the loss of plasma membrane proteins. Despite recent advances in artificial intelligence (AI) to infer cell types from multimodal data [27], conventional LCM-MS workflows remain limited in achieving cell-type resolution.

To overcome these limitations, we introduce *NicheProt*, a spatial proteomics platform that enables cell-type-resolved and tissue-compartment-specific proteomics with a photobleaching-mediated fluorescence cell barcoding method. Using 400 µm-thick and optically cleared tissue macrosections, we barcode target cells with a clear view of the surrounding three-dimensional (3D) tissue microenvironment with multiplex IF staining and confocal microscopy. Microscopic photobleaching of selected fluorescence signals creates unique optical barcodes for the target cell type located in defined regions of interest (ROIs). Following tissue dissociation, barcoded cells are isolated *via* fluorescence-activated cell sorting (FACS), preserving cellular integrity and protein content for subsequent MS analysis.

To validate our technique, we applied *NicheProt* to the mouse spleen, a key secondary lymphoid organ that is composed of distinct immune cell populations distributed across regional compartments, such as the T- and B-cell zones in the white pulp, red pulp, and marginal zone. While the splenic architecture is well-defined, the functional characteristics of individual immune cell types associated with their distinct compartments remain underexplored. In this study, we investigated CD11c^+^ dendritic cell (DC) responses to lipopolysaccharide (LPS)-induced inflammation. Using *NicheProt*, we isolated and profiled three distinct DC populations defined by their location inside or outside the T-cell zones in normal or inflamed spleen. Our results reveal distinct proteomic signatures between control and inflamed DCs, and between two spatially separated inflammatory DC subsets within the same LPS-treated spleen, identifying differences in 54 proteins. These compartment-specific profiles highlight the spatial regulation of DC function and identify potential biomarkers as well as candidates for novel therapeutics targeting specific cell populations.

## Results

### Development of NicheProt workflow

Current spatial proteomics technologies face challenges in providing comprehensive proteome profiling along with cell-type information. To address this limitation, we developed *NicheProt*, a spatial proteomics workflow that integrates five major steps: tissue cryopreservation and processing, photobleaching-mediated *in situ* fluorescence barcoding, tissue dissociation, fluorescence-activated cell sorting (FACS), and LC-MS/MS analysis (**Figure 1A**).

**Figure 1.**
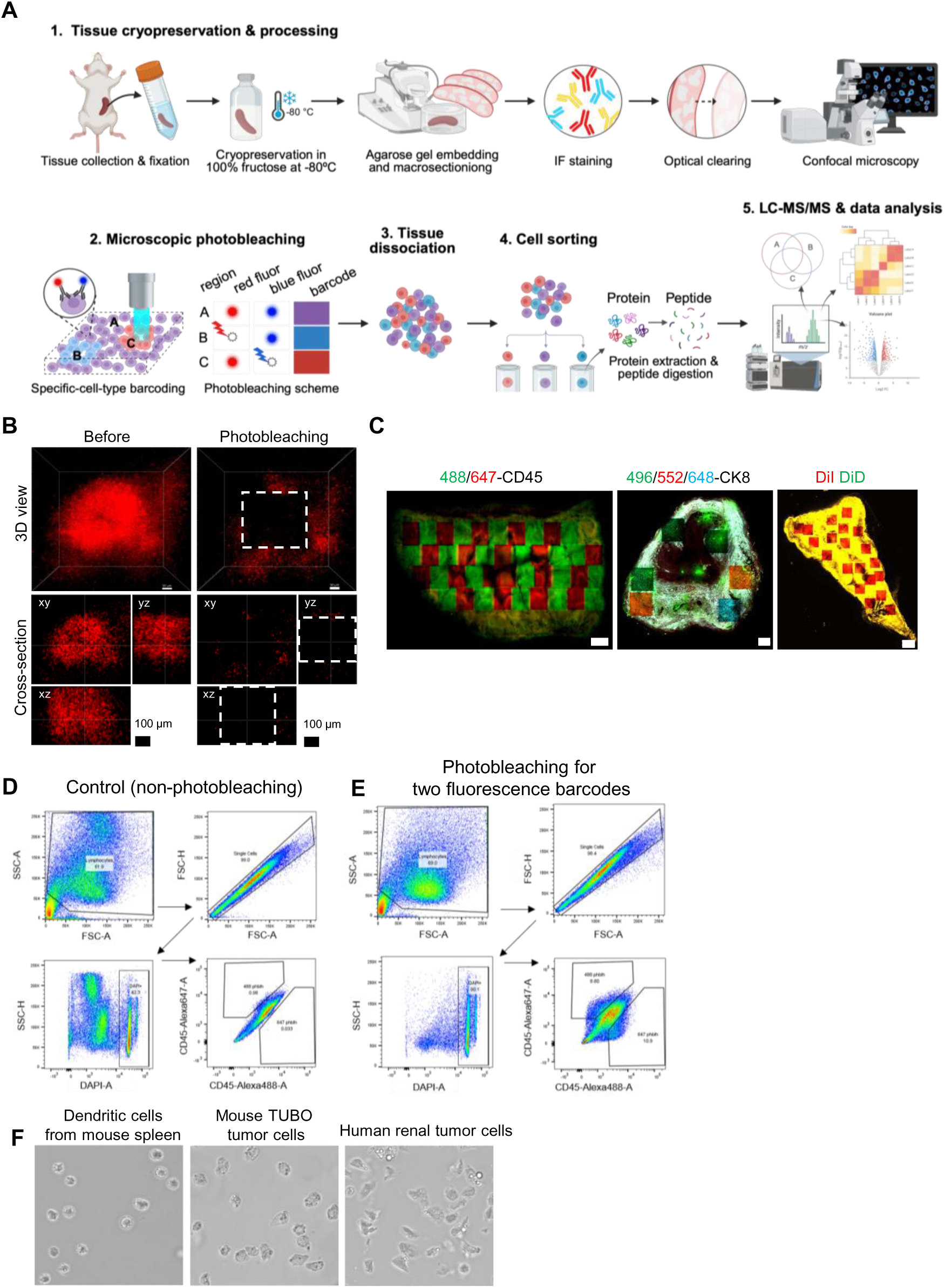
NicheProt workflow development. (A) NicheProt comprises five key steps in a sequential workflow: (1) tissue cryopreservation and processing, (2) microscopic photobleaching, (3) tissue dissociation, (4) cell sorting, and (5) LC-MS/MS and data analysis. (B) 3D rendering and virtual cross-sectional views of a photobleaching region in a 400 µm-thick mouse spleen macrosection stained with Alexa Fluor 647-anti-CD45 antibody. The photobleaching region was generated using a 640 nm excitation laser and a 20× air objective with a 3× zoom factor. Scale bar: 100 µm. (C) Photobleaching-mediated fluorescence barcoding in mouse tissues: (Left) two fluorescence barcodes in the mouse spleen targeting CD45^+^ lymphocytes. The spleen macrosection was stained with Alexa Fluor 488-conjugated (green) and Alexa Fluor 647-conjugated (red) anti-CD45 antibodies; (Middle) three fluorescence barcodes in the mouse ovarian tumor targeting PanCK^+^ tumor cells. The tumor macrosection was stained with Flamma 496-(green), Flamma 552-(red), and Flamma 648-(blue) anti-CK8 antibodies; (Right) a single-color fluorescence barcode in the mouse spleen stained with universal cell membrane dyes, DiI (red) and DiD (green). Scale bar: 500 µm. (D) Identification of the barcoded cells in the FACS plots. The control group consisted of dissociated splenocytes from the mouse spleen macrosection stained with DAPI, Alexa Fluor 488- and Alexa Fluor 647-anti-CD45 antibodies without photobleaching to establish the gating strategy. (E) The FACS plots show cells dissociated from the photobleaching mouse spleen macrosections (shown in **C** (Left)). Two fluorescence barcoded populations were identified by their intensity shift in the FACS plot relative to the non-photobleaching control in **D**. (F) Intact cell collection using the NicheProt approach. Various tissue samples, including mouse spleen, mouse mammary tumor, and human renal carcinoma, were tested to evaluate cell integrity post-cell sorting.

Building on our previous findings, we first optimized tissue fixation and cryopreservation protocols to preserve tissue status at the time of collection. Tissues were fixed in 4% paraformaldehyde (PFA) and then stored in 100% (w/v) D-fructose at -80 °C (**Figure 1A, step #1**). This approach effectively maintains overall tissue, cell, and protein integrity, enabling 3D confocal microscopy and intact cell collection in downstream steps. This preservation method supports a flexible processing timeline and facilitates sample sharing in multi-site studies, which is particularly important for ensuring robustness when handling clinical human specimens.

In the next step of the *NicheProt* workflow, tissues were thawed, washed in PBS, and embedded in agarose gel plugs for subsequent slicing into 400 μm-thick macrosections. Thick tissue macrosections retain 3D tissue architectures and minimize both cell disruption and protein loss. The macrosections were then stained with a cocktail of fluorophore-conjugated antibodies to visualize the distribution of a specific cell type. Tissue clearing is a chemical process that renders tissues optically transparent and is applied to facilitate deep microscopic visualization without disrupting light transmission. By combining D-fructose-based aqueous tissue clearing with multiplex IF microscopy [28, 29], we achieved 3D visualization of tissue architectures and identified specific cell types within defined ROIs in entire macrosections. To barcode a selective cell type, we used pairs of the same monoclonal antibodies conjugated to different fluorophores. For example, we used Alexa Fluor 488- and Alexa Fluor 647-anti-CD45 antibodies to stain CD45^+^ immune cells in the tissue macrosection (**Figure 1A, step #2**). Using a confocal microscope equipped with 405/488/561/640 nm excitation lasers, we photobleached one of the fluorophores to generate “optical barcodes”, enabling the *in situ* encoding of CD45^+^ immune cells with their positional information within three-dimensionally (3D) defined ROIs in the tissue macrosection (**Figure 1B and 1C, left**). Additional fluorescence barcodes could be produced by combining three antibodies conjugated to different fluorophores (**Figure 1C, middle**). Beyond targeting specific cell types, our photobleaching approach can also expand to universal cell barcoding in tissue macrosections stained with lipophilic membrane dyes such as DiI and DiD (**Figure 1C, right**). The volume of each photobleaching region was controlled by adjusting the objective magnification and zoom to precisely photobleach cells within the selected ROI (**Table S1**). Photobleaching each 3D region typically takes approximately 20 - 40 seconds using 20× and 40× air objectives with zoom factors, with higher magnifications and smaller regions requiring less time.

Following microscopic photobleaching, we isolated the barcoded cells from their tissue context. Spleen macrosections were mechanically dissociated, and barcoded cells were collected *via* FACS. These cells were identified by fluorescence intensity shifts in flow plots when comparing with “non-photobleaching” control cells dissociated from macrosections that did not undergo photobleaching-mediated barcoding (**Figure 1A, step #3-4**). Specifically, cells that were subjected to Alexa Fluor 647 bleaching exhibited a downward shift on the y-axis, while cells that were subjected to Alexa Fluor 488 bleaching showed a leftward shift on the x-axis in flow cytometry plots (**Figure 1D, E**). We employed a microfluidic-based mechanical cell sorter (MACSQuant® Tyto) for low-pressure cell sorting (< 3 psi) to ensure gentle handling and minimize the risk of cell rupture during the process. This approach enables the successful isolation of intact mouse splenocytes, mouse breast tumor cells, and human renal carcinoma cells from enzymatically or mechanically dissociated mouse and human tissues (**Figure 1F**). Finally, proteins were extracted, digested into peptides through validated methods, and analyzed by LC-MS/MS for bottom-up proteomic profiling (**Figure 1A, step #5**).

To evaluate cell and protein yields in the *NicheProt* workflow, we created 10 photobleached 3D regions using 20× and 40× objectives at different zoom settings in mouse spleen macrosections (**Table S1**). Because tissue cell density and protein content vary across different tissues and cell types, a preliminary assessment of cell and protein recovery from each target tissue and cell type is essential for designing a practical *NicheProt* analysis plan tailored to specific research goals. This information guides the determination of the number and volume of photobleached regions required per tissue macrosection to ensure sufficient cell and protein yields for subsequent standard LC-MS/MS analysis.

### Protein artifact assessment during NicheProt processing

Given the multi-step nature of our spatial proteomics workflow, we first evaluated whether tissue cryopreservation introduces protein artifacts or not. We compared protein profiles from mouse spleens processed immediately after fixation with those cryopreserved in 100% D-fructose at -80 °C prior to extraction (**Figure 2A**). Our analysis revealed consistency in the protein profiles following D-fructose cryopreservation. We quantified up to 4,818 proteins per sample with a sample-by-sample Pearson correlation ranging from 0.964 to 0.985 and a strong correlation of 0.994 between control and cryopreserved groups (**Figure 2B-D**). In the PCA plot, samples from both groups were intermingled without clear separation, indicating remarkable similarity in their overall features (**Figure 2E**). Volcano plot analysis showed no significantly altered proteins, and coefficients of variation (CVs) were below 0.2 in both conditions, indicating that the proteomics data was highly reproducible across biological replicates (**Figure S1A-C**). These results validate that cryopreservation does not introduce detectable protein artifacts and enhances workflow flexibility by allowing long-term storage, simplified sample sharing for collaborative studies, and archiving clinical human tissue samples.

**Figure 2.**
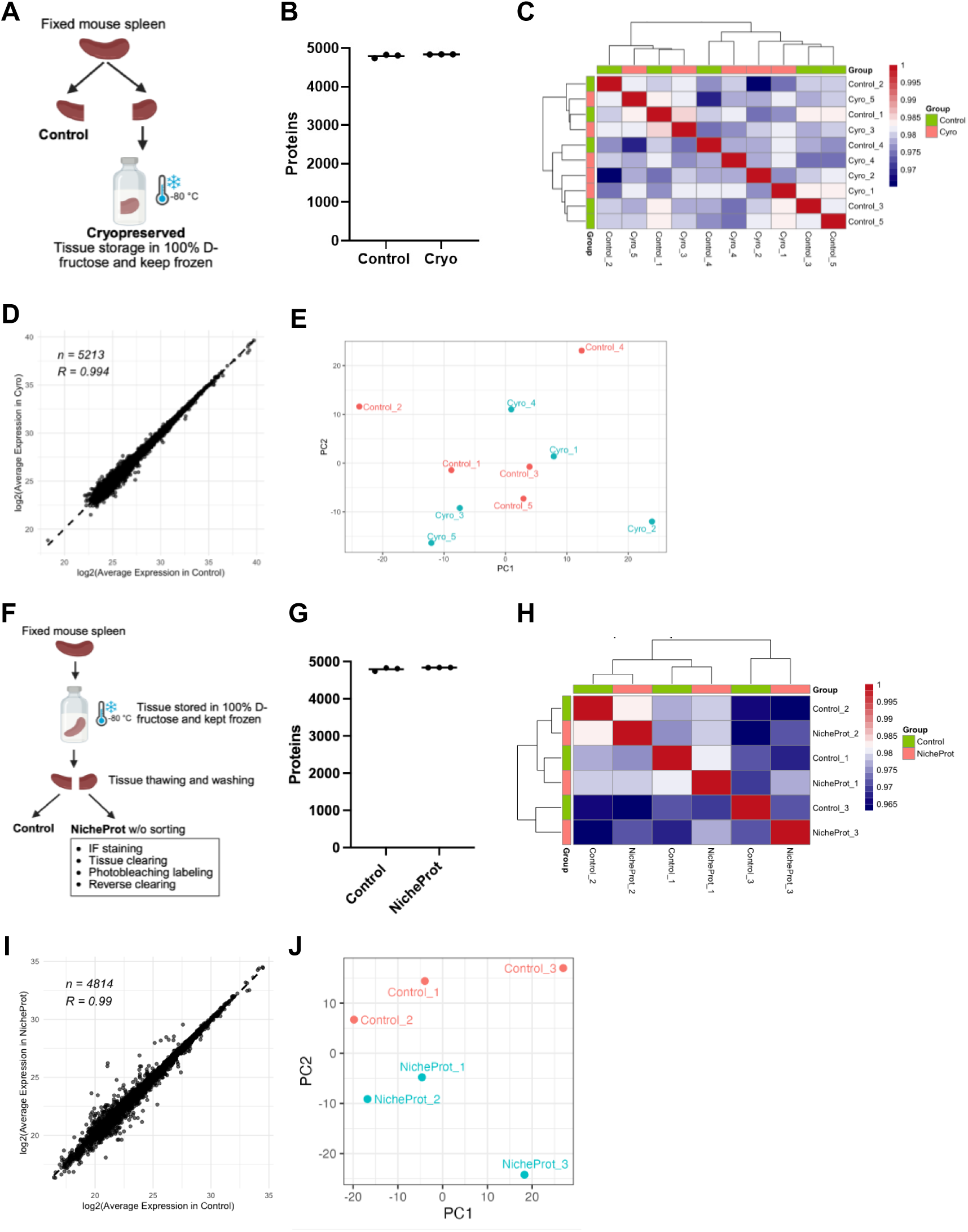
Proteome artifact evaluation during the NicheProt procedure. (A) Experimental design comparing protein expression between control and cryopreserved spleens. (B) The number of protein identifications between the control and cryopreservation groups (n=5 per group) obtained using LC-MS/MS. (C) Pearson correlation matrix showing sample-to-sample comparison. A high correlation indicates a similar proteome profile between control and cryopreserved samples. (D) The correlation analysis by groups with the correlation coefficient (R) equal to 0.994 demonstrates remarkable similarity between the control and cryopreservation groups. (E) Principal component analysis (PCA) of the control and cryopreserved samples. (F) Experimental design comparing protein expression between control and NicheProt-processed spleens. (G) The number of protein identifications between the control and NicheProt groups (n=3). (H) Pearson correlations of sample-to-sample comparison between control and NicheProt samples. (I) The correlation analysis by groups with the correlation coefficient (R) equal to 0.99. (J) PCA of the control and NicheProt groups.

Next, we tested whether the *NicheProt* workflow itself affects protein expression. Using cryopreserved mouse spleens, we compared protein profiles between non-processed controls and *NicheProt*-processed tissues (**Figure 2F**). The *NicheProt*-processed tissues were stained with a cocktail containing DAPI, Alexa Fluor 488-anti-CD45, and Alexa Fluor 647-anti-CD45 antibodies, followed by photobleaching of Alexa Fluor 647 in arbitrary regions to generate the fluorescence barcodes on CD45^+^ immune cells in the spleen macrosection. Optical tissue clearing and reverse clearing (from D-fructose to PBS) were performed before and after photobleaching, respectively. Since the control group was not stained with a fluorescent antibody, the FACS step was excluded from this comparison to avoid potential bias arising from cell type selection in the *NicheProt*-processed group. Previous studies have shown that microfluidic cell sorting does not significantly affect protein profiles due to its gentle and low-air pressure (<3 psi) mechanism [30, 31]. We detected close to 5,000 proteins per sample with Pearson correlations ranging from 0.965 to 1.000 across conditions and an intergroup correlation of 0.99 (**Figure 2G-I**). In the PCA plot, PC1 accounts for 95% of the variance, while PC2 represents less than 5%. Sample separation along PC1 was driven by biological variability among mice rather than by the *NicheProt* processing (**Figure 2J**). Additionally, no significantly up- or downregulated proteins were observed between the groups in the volcano plot, and CVs remained below 0.2, confirming high reproducibility (**Figure S1D-F**). Together, these findings validate *NicheProt* as a reliable spatial proteomics workflow that preserves protein integrity and avoids significant artifacts.

### Proteomic profiling of dendritic cell responses to LPS in mouse spleens

The spleen is the largest secondary lymphoid organ in the body and plays a vital role in orchestrating innate and adaptive immune responses in mice and humans [32, 33]. The rodent spleen comprises the red pulp (RP), white pulp (WP), and marginal zone (MZ), which lies between these two regions [34, 35]. The WP serves as the main immunological hub and is divided into B and T cell zones. Upon encountering antigens or pathogens, DCs in the spleen act as key antigen-presenting cells (APCs) that initiate T cell activation and the subsequent adaptive immune response [36, 37]. During inflammation, DC migration into the T cell zone (TCZ) is essential for T cell priming [34, 38]. Previous studies have shown that migratory DCs undergo significant proteomic remodeling, acquiring more mature, stimulatory, and cytokine-secreting phenotypes compared with naïve DCs [39–41]. However, spatially associated subtypes and functional roles of migratory DCs in response to immune stimuli require further elucidation.

In the normal mouse spleen, most DCs reside in the MZ, outside the TCZ. In contrast, multiplex IF confocal microscopy revealed CD11c⁺ DC migration toward the TCZ after lipopolysaccharide (LPS) administration, a gram-negative bacterial cell wall component widely used to induce systemic acute inflammation in mice [42–45] (**Figure S2A**). Furthermore, we identified two spatially distinct DC populations in the LPS-treated mouse spleen: one that migrated into the TCZ and another that remained outside (**Figure S2B**). In this study, we applied *NicheProt* to characterize DC subsets and their immunoregulatory functions in distinct tissue compartments. We barcoded and isolated three CD11c^+^ DC groups: (1) DCs located outside the TCZ in the normal spleen (“*norm*”), (2) DCs located outside the TCZ in the LPS-treated spleen (“*outside*”), and (3) DCs that had migrated inside the TCZ in the LPS-treated spleen (“*inside*”) (**Figure 3A**). Spleen macrosections were stained with Alexa Fluor 488- and Alexa Fluor 647-conjugated anti-CD11c antibodies to label DCs, and DyLight 550-conjugated anti-CD3 antibody to mark the TCZ. For example, to barcode DCs *inside*, we photobleached Alexa Fluor 647-anti-CD11c in the TCZ, converting the pseudocolor of DCs from yellow to green (**Figure 3B, C**). We photobleached multiple ROIs to ensure sufficient cell number and protein yield collected from the target compartment. From a single 400 µm-thick LPS-treated spleen macrosection, we isolated 8.3 ×10^5^ DCs, yielding 22.37 µg of protein for LC-MS/MS analysis (**Figure 3D**). Similar procedures were followed to isolate DCs in *norm* and *outside* groups from their respective compartments in the normal and LPS-treated spleens (**Figure 3E, S3**).

**Figure 3.**
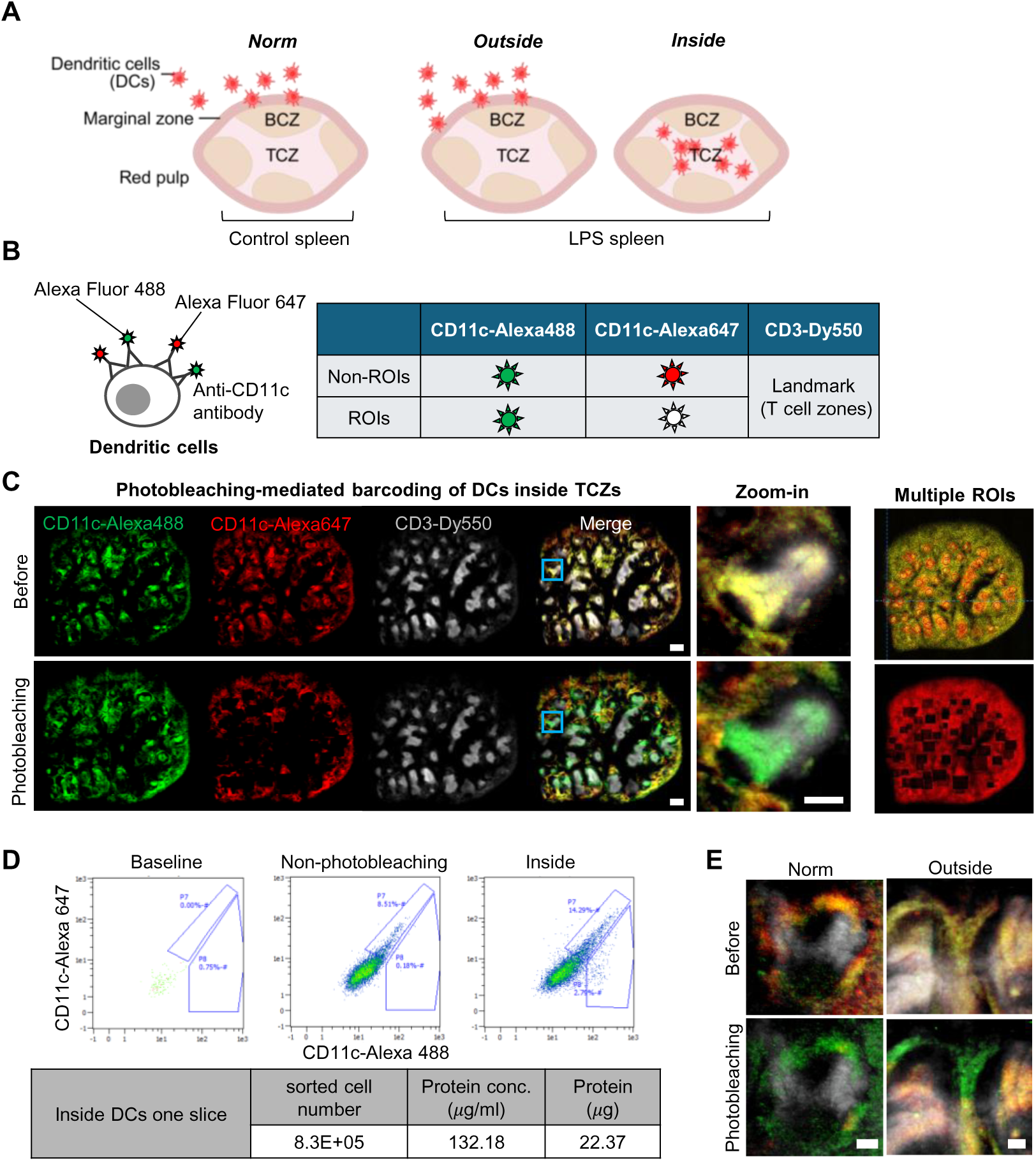
DC barcoding in mouse spleen compartments by microscopic photobleaching. (A) Three DC populations, *norm*, *outside*, and *inside* from normal and LPS-treated mouse spleens, were analyzed using NicheProt. (B) DCs were stained with Alexa Fluor 488 and Alexa Fluor 647-anti-CD11c antibodies for visualization and photobleaching. DCs in multiple ROIs were barcoded by photobleaching the Alexa Fluor 647 signal, creating a distinct fluorescence signature from the rest of the DCs in non-photobleaching regions. (C) Example of photobleaching barcoding on DCs in the *inside* group. Photobleaching of Alexa Fluor 647 on CD11c⁺ DCs converted yellow signals (Alexa Fluor 488 and 647) to green (Alexa Fluor 488) within multiple CD3⁺ TCZs. The zoom-in image of the marked area shows a high-resolution TCZ and precise photobleaching of the DCs. (Right) Multiple ROIs were photobleached and collected from a single half-spleen slice. Scale bars: 500, 200 µm. (D) FACS plots show the fluorescence intensity shift of Alexa Fluor 647 in barcoded DCs compared to the signals in the “baseline” and “non-photobleaching” control groups. The table below summarizes the cell number, protein concentration, and protein amount obtained from the barcoded DCs. (E) High-resolution microscope images of DCs in the *norm* and *outside* groups before and after photobleaching. Scale bars: 100 µm. BCZ: B cell zone; TCZ: T cell zone.

Across biological replicates, we quantified 4,741 proteins and observed a clear separation among the three DC subsets *via* PCA and heatmap clustering. (**Figure 4A, B**). Several inflammatory proteins, including aconitate decarboxylase 1 (Acod1), radical S-adenosyl methionine domain-containing 2 (Rsad2), interferon-induced protein with tetratricopeptide repeats 3 (Ifit3), 2’-5’ oligoadenylate synthetase 1A (Oas1a), and guanylate binding protein 5 (Gbp5), were significantly upregulated in both DCs from the *inside* and *outside* groups, consistent with known responses to LPS [46–48]. Additionally, S100 family proteins, C-X-C motif chemokine ligand 10 (Cxcl10), and interleukin 4 induced 1 (Il4i1) were markedly elevated in DCs from the *inside* group compared to the *norm* group (**Figure 4C**). These proteins are known for their proinflammatory roles during inflammation. S100 calcium-binding protein A8 (S100A8) and S100 calcium-binding protein A9 (S100A9) are critical calcium-binding proteins that promote immune cell recruitment and cytokine production [49, 50]; Cxcl10 functions as a chemokine that recruits leukocytes and enhances cytotoxic T cell activation [51, 52]; and Il4i1, predominantly expressed in antigen-presenting cells like DCs and macrophages, plays an immunoregulatory role during inflammation [53–55]. Likewise, numerous inflammatory proteins were differentially expressed in DCs *outside* in comparison to DCs *norm* (**Figure 4D**). Gene ontology (GO) enrichment analysis revealed that DCs from both *inside* and *outside* groups were enriched for pathways related to type I and II interferon responses, cytokine production, innate immune activation, bacterial defense, and LPS response (**Figure 4E, F**). These results demonstrate successful validation of our *NicheProt* analysis in agreement with known reference data reported in multiple studies.

**Figure 4.**
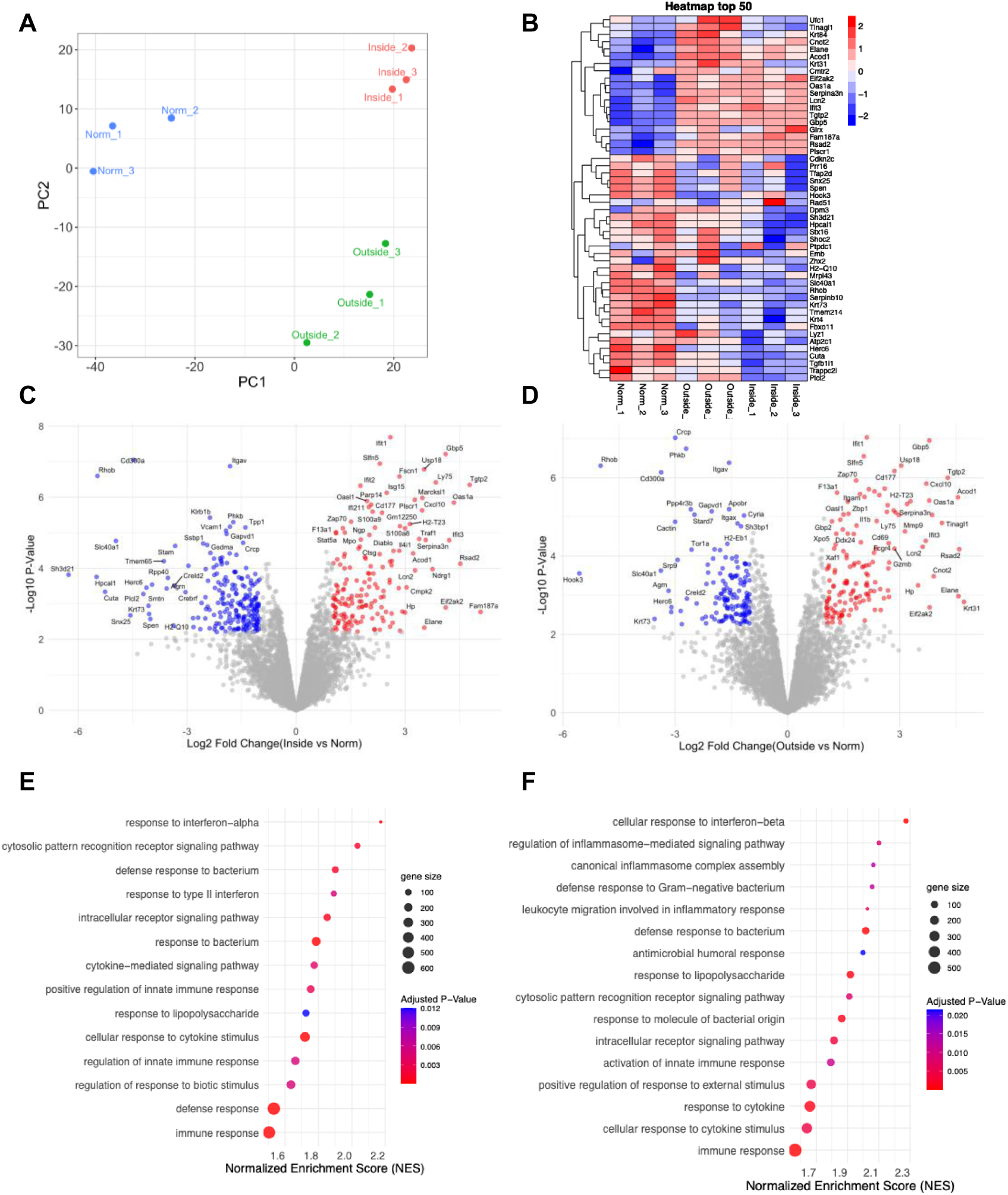
NicheProt reveals distinct protein profiles between DC populations. (A) PCA plot of the proteome collected from DCs in the *norm*, *outside*, and *inside* groups (n=3). (B) Heatmap of the top 50 most variable proteins. (C) Differential protein expression of DC *inside* versus *norm* groups. Proteins upregulated in the DC *inside* group are highlighted in red, while those upregulated in the DC *norm* group are shown in blue. (D) Differential protein expression of DC *outside* versus *norm* groups. Proteins upregulated in the DC *outside* group are highlighted in red, while those upregulated in the DC *norm* group are shown in blue. (E, F) Gene ontology (GO) enrichment pathway analysis of biological pathways differentially regulated (E) between DC *inside* and *norm* groups and (F) between DC *outside* and *norm* groups.

### Spatially distinct dendritic cell populations in the inflamed mouse spleen

In the LPS-treated spleen, we observed two spatially distinct DC populations: one that migrated into the TCZ (the *inside* group) and another that remained outside the TCZ (the *outside* group). As no previous study has directly compared these subsets, we utilized *NicheProt* to profile their whole proteomes. Differential expression analysis revealed 54 proteins with significantly altered abundance between the two populations (**Figure 5A**). GO pathway analysis indicated that DCs from the *inside* group were enriched for aggrephagy, a cellular mechanism for clearing protein aggregates to maintain homeostasis and limit excessive inflammation. In contrast, DCs from the *outside* group were enriched for pathways related to intermediate filament organization, suggesting increased surface receptor turnover likely due to active endocytosis or antigen patrolling (**Figure 5B**). These findings reveal proteomic differences between compartment-specific DCs that could lead to uncharacterized DC subtypes with distinct functional roles in the spleen microenvironment.

**Figure 5.**
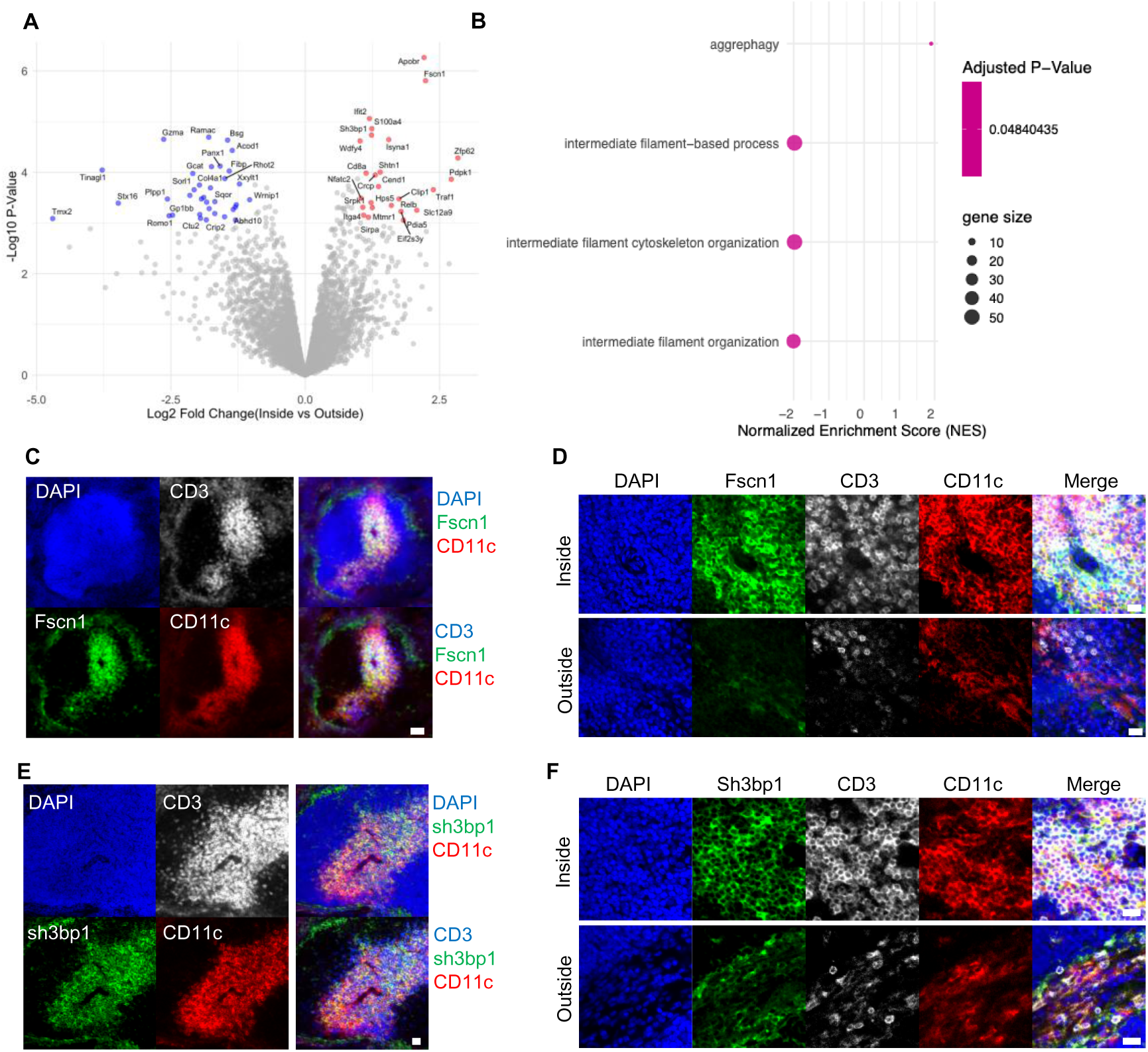
Spatially distinctive DC populations in LPS-treated mouse spleen. (A) Differential protein expression between the DC *inside* versus *outside* groups. Proteins upregulated in the DC *inside* group are highlighted in red, while those upregulated in the DC *outside* group are shown in blue. (B) GO enrichment pathway analysis of biological pathways differentially regulated between the DC *inside* versus *outside* groups. (C-F) Multiplex IF images confirmed increased (C, D) Fscn1 and (E, F) Sh3bp1 expression of the DCs *inside* the TCZ in lower and higher-resolution images. Scale bars: 50, 20 µm.

These 54 differentially expressed proteins may serve as potential biomarkers for distinct DC populations as well as possible protein targets for novel therapies. Among the proteins upregulated in *outside* DCs, we identified granzyme A (GzmA), granzyme B (GzmB), platelet endothelial cell adhesion molecule 1 (Pecam1, CD31), Acod1, integrin alpha-D (Itgad), normal mucosa of esophagus-specific gene 1 protein (Nmes1), and tubulointerstitial nephritis antigen-like 1 (Tinagl1). GzmA, expressed in plasmacytoid DCs, promotes pro-IL-1β production and enhances cross-priming of conventional DCs [56, 57]. GzmB is implicated in immune regulation and extracellular matrix (ECM) remodeling. Pecam1 has been associated with reduced proinflammatory DC maturation [58]. Acod1 modulates immune tolerance to LPS, suppressing overactive responses while supporting antimicrobial activity [59–61]. While Itgad (encoding CD11d) is primarily expressed on monocytes and macrophages, its presence on DCs may facilitate ECM adhesion and migration to inflammatory sites [62, 63]. Nmes1 is recognized as a marker of effector DCs and is strongly upregulated in LPS-treated DCs [64]. Though understudied in DCs, Tinagl1 has been reported to suppress tumor progression in breast cancer models, suggesting a possible regulatory role in inflammation that warrants further investigation [65].

In contrast, proteins significantly upregulate in *inside* DCs included apolipoprotein B receptor (Apobr), S100 calcium-binding protein A4 (S100a4), interferon-induced protein with tetratricopeptide repeats 2 (Ifit2), T-cell surface glycoprotein CD8 alpha chain (Cd8a), ITGA4 integrin subunit alpha 4 (Itga4), fascin actin-bundling protein 1 (Fscn1), and SH3 domain binding protein 1 (Sh3bp1). Apobr facilitates the uptake of lipoproteins, such as low-density (LDL) and high-density (HDL) lipoproteins, which may aid in LPS clearance during bacterial inflammation [66–68]. S100a4 promotes cytokine production and immune cell recruitment and is linked to inflammatory diseases such as rheumatoid arthritis and systemic sclerosis [69, 70]. Ifit2 is a key mediator of type I interferon (IFN-1) signaling, drives tumor necrosis factor-alpha (TNF-α) or interleukin 6 (IL-6) production, and is highly responsive to LPS stimulation. Inhibition of Ifit2 reduces inflammatory cytokine levels and mortality in the endotoxin shock model [71]. CD8a is a surface marker of a specific DC subset and functions as a co-receptor for antigen cross-presentation on MHC class I to CD8^+^ T cells [72]. Itga4 is involved in adhesion and migration by interacting with fibronectin and VCAM1, marking DC activation and trafficking to inflamed sites [73–75].

To validate our findings with an orthogonal approach, we performed IF staining for two upregulated proteins found in DCs from the *inside* group, Fscn1 and Sh3bp. Fscn1 is an actin-bundling protein that is highly induced during DC maturation and is required for effective membrane protrusion and migration. It also serves as a biomarker of effector DCs during allergen responses [64]. Sh3bp1 is involved in cytoskeletal reorganization and morphological adaptation during cell migration [76–78]. Our IF results confirmed higher expression of both proteins in the *inside* DCs compared to *outside* DCs, consistent with the *NicheProt* results (**Figure 5C-F**).

## Discussion

LCM coupled with MS is an established technology that enables spatially resolved proteomics analysis of tissue subregions. However, LCM-MS has limited ability to provide accurate cell-type specific proteome information because these dissected subregions often contain heterogeneous cell populations and other non-cellular components (e.g., extracellular matrix (ECM)). In our previous attempt to isolate individual cells from FFPE thin tissue sections using a laser microdissection system, we observed frequently damaged cell membranes, leading to protein loss and the need to collect large numbers of cells. In addition, the whole LCM process could be time-consuming and labor-intensive. These challenges highlight the unmet need for a new method that enables the selective and efficient collection of specific cell types from defined tissue compartments for whole-proteome analysis.

We here demonstrate *NicheProt*, a spatial proteomics approach that leverages microscopic photobleaching to optically encode positional information on specific cell types within selected ROIs, thus avoiding the need for genetically engineered mouse models expressing fluorescence proteins [79]. This capability makes *NicheProt* adaptable to a broad range of tissue samples, both in the preclinical and clinical settings. Designed for general laboratory use, *NicheProt* is cost-effective and compatible with common reagents, commercially available fluorophore-conjugated antibodies, and standard lab and core facility equipment, unlike many other spatial omics platforms that require specialized instrumentation and costly supplies. In *NicheProt*, optical clearing and multiplex IF confocal microscopy of tissue macrosections provide 3D visualization of the tissue microenvironment, enabling precise ROI selection for photobleaching labeling. Using antibody staining, we identified and labeled specific cell populations for collection while preserving their integrity by working on thick tissue slices to avoid cutting through cells. Importantly, IF staining with fluorescent antibodies in *NicheProt* is not used for direct detection of proteins in proteome analysis. Instead, we performed unbiased, bottom-up proteome profiling using LC-MS/MS, identifying up to 5,000 proteins per sample collection. *NicheProt* substantially expands protein analytical coverage beyond what is achievable with antibody-based detection methods.

We validated that the *NicheProt* workflow does not introduce significant artifacts to the protein profiling results (**Figure 2 and Figure S1**). Moreover, our tissue cryopreservation protocol, using PFA fixation followed by incubation and storage in 100% D-fructose at -80 °C, offers flexibility to experimental scheduling and facilitates sample sharing and reproducibility across research sites. Finally, NicheProt captures proteomic information of specific cell types within their niches, revealing how microenvironmental cues and intercellular interactions shape their unique roles in the functional tissue compartment for tissue homeostasis and disease pathogenesis. We validated the feasibility of the *NicheProt* method by comparing the proteome of CD11c^+^ DCs located either inside or outside the TCZs of LPS-treated mouse spleen to those in normal mouse spleens. Our results were consistent with the reported protein signatures of LPS-induced immune activation, supporting the reliability of the *NicheProt* analysis. Gene ontology (GO) enrichment analysis revealed key pathways associated with cytokine production, interferon responses, innate immune activation, and, most notably, response to LPS, reflecting the expected molecular changes in DCs during inflammatory conditions. Several well-characterized inflammatory proteins, including Rsad2, Ifit3, Oas1a, Gbp5, and Acod1, were significantly upregulated, as shown in the heatmap and differential expression analysis (**Figure 4C, D**), consistent with the previously published studies [46–48]. Rsad2 plays a crucial role in DC maturation and immune activation *via* Toll-like receptor (TLRs) 7 and 9 signaling [80, 81]; Interferon-induced proteins with tetratricopeptide repeats (IFITs) are robustly induced by LPS. Specially, Ifit3 promotes innate immune response against bacterial infections *via* type I interferon (IFN) signaling [82, 83]; Oas1a is another type I IFN-responsive protein that activates the innate immune system against bacterial infections [84, 85]; Gbp5 participates in type II interferon signaling and inflammasome complex assembly [86, 87]; Acod1 is more complex since it is highly induced by LPS in macrophages and DCs and has both pro- and anti-inflammatory roles. While Acod1 supports antimicrobial defense, it also contributes to immune tolerance by attenuating excessive innate responses [59–61]. Further mechanistic studies are needed to fully understand the dual functions of Acod1. Additional inflammatory markers, such as S100 family proteins, Cxcl10, and Il4i1, were also significantly upregulated in DCs from both *inside* and *outside* groups, further supporting the findings in our spatial proteomics results.

When directly comparing the two spatially distinct DC populations within the LPS-treated mouse spleen, which have not been previously studied, we identified 54 differentially expressed proteins (**Figure 5**). Among them, Apobr, S100a4, Ifit2, Cd8a, Itga4, Fscn1, and Sh3bp1 were significantly upregulated in the DCs from the *inside* group, while GzmA, GzmB, Pecam1, Acod1, Itgad, Nmes1, and Tinagl1 were upregulated considerably in the DCs from the *outside* group. To validate our proteomics findings, we performed multiplex IF confocal microscopy for Fscn1 and Sh3bp1, two proteins that regulate DC maturation and migration [88, 89] and were enriched in DCs from the *inside* group. The imaging results confirmed higher expression of both proteins in DCs from the *inside* group compared to those from the *outside* group, consistent with the *NicheProt* results. Interestingly, GO enrichment analysis did not reveal strong pathway-specific associations that could clearly define either subset as strictly proinflammatory, anti-inflammatory, migratory, or stationary. The DCs in the *inside* and *outsid*e groups may represent closely related functional states rather than distinct phenotypes. This result demonstrates that *NicheProt* can resolve the spatial and functional dynamics of a specific cell type by detecting subtle protein-level changes during inflammation. Further mechanistic studies will be needed to elucidate the detailed roles of these DC subsets in immune responses. The 54 differentially expressed proteins identified in this study represent may serve as therapeutic targets for inflammatory diseases. For instance, Ifit2, which was significantly upregulated in the DC *inside* group, has been strongly associated with gram-negative septic shock. Previous studies have demonstrated that Ifit2-deficient mice exhibit significantly reduced serum levels of IL-6 and TNF-α and improved survival in endotoxin shock models [71]. These findings suggest that inhibiting Ifit2 using small molecules, such as glucocorticoids [90], may offer a viable therapeutic strategy for treating septic shock.

In summary, *NicheProt* enables the detection of proteomic differences among the same cell type residing in distinct functional tissue microenvironments. Its sensitivity allows the resolution of protein detection of cellular subsets to as few as a handful of differentially expressed proteins. This platform holds great potential for uncovering previously unrecognized cell subtypes and elucidating their roles within the spatial tissue context in regulating homeostasis and pathogenesis.

## Limitations and future directions

Several additional optimizations could further advance the *NicheProt* workflow in the future, including automated microscopic photobleaching, an increased number of fluorescence barcodes per tissue slice, and enhanced spatial resolution. In the current prototype, photobleaching is performed manually by adjusting the microscope stage to target different ROIs. A key upgrade would be the integration of an automated stage and software system, allowing users to predefine imaging routes, photobleaching sequences, laser exposure time, and other microscope settings. This automation would significantly increase throughput for barcoding specific cell types in multiple ROIs and make the workflow more accessible for new users.

Secondly, the number of fluorescence barcodes per tissue macrosection is currently restricted by four available excitation lasers (405, 488, 561, and 640 nm) on our confocal microscope. When one channel is reserved for a tissue compartment marker (e.g., CD3 for marking the TCZs in the mouse spleen), the remaining three lasers can generate up to six binary (1 or 0) fluorescence barcodes *via* complete photobleaching. To expand the number of barcodes beyond this, gradient photobleaching to varying degrees through adjusting laser power and exposure time might be achievable [91]. Accurate discrimination of these gradient-barcoded cell populations will require optimization of the FACS setup, particularly its sensitivity and gating strategies for detecting varying fluorescence intensities.

In addition, the spatial resolution of the current version of *NicheProt* is determined and affected by the cell loss during tissue dissociation, the sorting efficiency, and the LC-MS/MS sensitivity. To mitigate this limitation, we photobleached multiple areas within the same tissue compartment to ensure adequate recovery of the target cell population. Achieving resolution from a single photobleaching area will require advanced cell separation techniques and ultra-sensitive MS [92–94] to minimize loss of barcoded cells during the multi-step workflow and to enable MS sequencing from a smaller starting sample amount. This enhancement will allow proteomic analysis of rare cell populations in a smaller tissue compartment.

We demonstrated the feasibility of *NicheProt* using the mouse spleen as a tissue model due to its accessibility and compatibility with mechanical dissociation, which eliminates the need for enzymatic digestion and minimizes variability during early-stage method development. However, extending *NicheProt* to other tissues, such as tumors, brain, and kidney, will require further optimization of enzymatic dissociation protocols and, in some cases, protein (or antigen) retrieval steps to enhance protein detectability in heavily fixed tissue samples.

Future expansions of the *NicheProt* platform could incorporate additional spatial omics methods. For instance, combining *NicheProt* with spatial transcriptomics technologies (e.g., Visium/Xenium by 10x Genomics), cyclic multiplex IF imaging platform, or computational image analysis tools [22, 24, 95, 96] could provide further information on the tissue microenvironment to guide ROI selection for photobleaching-barcoding of the target cell populations. This integration is particularly valuable for tumor specimens, which often lack the well-defined anatomical compartments seen in normal tissues. Furthermore, AI-based data integration of single-cell sequencing or flow cytometry datasets using advanced computational methods (e.g. SCPro [97]) would support deeper cellular phenotyping. These enhancements would further advance the capabilities of using *NicheProt* for discovering novel cell subtypes and therapeutic targets.

## Methods

### Inflammation mouse model

Normal and inflamed spleens were harvested from 6 to 8-week-old male C57BL/6J mice. Lipopolysaccharide (LPS) purified from Escherichia coli 0111:B4 (Cat# LPS25, Sigma-Aldrich) was reconstituted in ddH_2_O to a concentration of 5 mg/ml, and the aliquots were stored at -20 °C until use. The working concentration of LPS was diluted to 0.2 µg/µl on the day of the experiment. Mice were intravenously injected with 15 µg of LPS in the tail vein. After 6 hours, LPS-injected mice were sacrificed, and spleens were collected. Normal spleens were obtained from healthy mice without any treatment. All animal procedures were performed in accordance with an approved protocol (ACC# 24-107) from the Institutional Animal Care and Use Committee at the University of Illinois Chicago.

### Tissue cryopreservation and macrosectioning

Spleen tissues were harvested from healthy 6 to 8-week-old male C57BL/6J mice without any treatment. Immediately after collection, the tissues were transferred to a 4% paraformaldehyde (PFA) solution prepared in PBS and fixed for 20 minutes at room temperature (RT). Following fixation, tissues were washed three times with cold PBS at 4 °C for 5 minutes each. These spleens were either used immediately or cryopreserved. For cryopreservation, spleens were incubated in 100% (w/v) D-fructose (Cat# F0127, Sigma-Aldrich) solution prepared in 100 mM phosphate buffer (PB) (pH 7.8) with gentle agitation for 1 hour at RT to allow equilibration and then transferred to a -80 °C freezer. On the day of the experiment, frozen spleens were thawed and washed four times with cold PBS at 4 °C for 5 minutes each to remove residual fructose. The spleens were then embedded in 2% agarose gel (LE Quick Dissolve Agarose, GeneMate) dissolved in distilled water and mounted on a vibratome (VT1200S, Leica) once the gel had solidified. The vibratome chamber was filled with cold PB and surrounded by crushed ice in a designated tray to maintain a cold environment. Spleen tissue was serially sectioned into 400 µm-thick macrosections.

### Immunofluorescence (IF) staining

Monoclonal purified anti-mouse CD3 and CD45 antibodies were purchased from BioLegend and conjugated with fluorophores *via* N-hydroxysuccinimide (NHS)-ester chemistry. Other antibodies were purchased pre-conjugated with fluorophores from BioLegend. Detailed information on antibodies and fluorophores is provided in **Table S2**. Fluorophore conjugation was performed by incubating a designated ratio of dye to purified antibody overnight at 4 °C under gentle agitation. Unconjugated dye molecules were removed through dialysis in PBS using cassettes (MWCO 10K, Cat# 66383, ThermoFisher Scientific) at 4 °C over three days. PBS was replaced three times during the dialysis process. The final conjugated antibody was transferred to clean microcentrifuge tubes and stored at 4 °C. Staining buffer was prepared by dissolving bovine serum albumin (BSA) (Cat# F0127, Sigma-Aldrich) in RPMI 1640 medium (Cat# 22400089, Gibco, ThermoFisher) at 10 mg/ml. Spleen macrosections were stained with antibody cocktails in the staining buffer for 20 hours at 4 °C under gentle agitation. After the staining, the macrosections were washed three times in cold PBS at 4 °C for 5 minutes each.

To validate protein upregulation identified in *NicheProt* using IF staining, the LPS-treated spleen macrosections were stained with primary anti-Fscn1 (3:500 vol/vol) (Cat# 14384-1-AP, Proteintech, ThermoFisher Scientific) or anti-Sh3bp1 antibodies (3:500 vol/vol) (Cat# 20541-1-AP, Proteintech, ThermoFisher Scientific) in the staining buffer for 18 hours at 4 °C under gentle agitation. Afterwards, spleen macrosections were washed three times in cold PBS at 4 °C for 5 minutes each and stained with secondary Alexa Fluor 488-conjugated goat-anti-rabbit antibody (Cat# 111547003, Jackson ImmunoResearch Laboratories, 2.4 mg/ml) at a 1:100 vol/vol dilution ratio in the staining buffer for 18 hours at 4 °C under gentle agitation. Spleen macrosections were washed again with cold PBS before tissue clearing and microscopy.

### Tissue optical clearing and reverse

D-fructose (Cat# F0127, Sigma-Aldrich) solutions were prepared in 100 mM PB buffer (pH 7.8) and used as aqueous-based tissue clearing reagents. Following IF staining, spleen tissue macrosections were sequentially incubated in 50% (w/v) D-fructose solution for 20 minutes and then in 80% (w/v) D-fructose solution for 30 minutes at RT. After photobleaching labeling on the tissue, the cleared tissue macrosections were subjected to a reverse tissue clearing process by incubating them in 50% (w/v) D-fructose solution for 20 minutes at RT, followed by four washes in cold PBS at 4 °C for 5 minutes each. This step gradually reversed the tissue status to its original condition.

### Microscopic photobleaching barcoding on cells

Every cleared spleen macrosection in 80% (w/v) D-fructose solution was placed on a microscope slide (Cat# 125442, Fisher Scientific) and covered with a coverslip (Cat# 12541032, Fisher Scientific) for imaging and photobleaching barcoding. Photobleaching was performed using a Caliber I.D. RS-G4 confocal microscope. The laser channels were calibrated internally to ensure an optimal laser output of 100% before each experiment. The maximum operating laser power of both 488 nm and 561 nm channels is 100 mW, while for the 640 nm channel is 70 mW. To maintain the cold condition during the photobleaching, dry ice was placed on a tray near the microscope stage, and a fan was blowing cool air toward the imaging tissue. For an initial scanning, we used a 10× objective (Olympus UPLXAPO 10×, NA: 0.4, WD: 3.1 mm) to visualize the overall tissue landscape to define ROIs and select target regions for barcoding. A 100 μm z-stack image was captured as a representative pre-photobleaching image. The objective was then switched to a 20× air objective (Olympus UPLXAPO 20×, NA: 0.8, WD: 0.6 mm) to reassure the 3D tissue microenvironment within ROIs. Only regions without compartmental ambiguity were selected for photobleaching barcoding. For barcoding, either the 488 nm or 640 nm laser was set to 95% intensity output and applied for 30-40 seconds to photobleach Alexa Fluor 488 or Alexa Fluor 647 signal, respectively. Zoom factors were adjusted to precisely confine the photobleaching regions within each selected ROI. Multiple photobleachings were generated within a single tissue macrosection to label cells from the same tissue niche. Finally, a post-photobleaching 100 μm z-stack image was acquired using a 10× objective to demonstrate the barcoded regions in each tissue slice.

### Spleen dissociation

Two to three mouse spleen macrosections were transferred to an EASYstrainer with a 40 μm mesh size (Cat# 07-001-106, Fisher Scientific) positioned atop a 1.5 ml tube. The macrosections were gently minced against the mesh filter using a 1 ml syringe plunger in circular motions. Dissociated splenocytes were washed through the mesh filter by adding 600-800 μl of cold PBS. Additional mincing and PBS were applied if undissociated spleen tissue fragments remained. The resulting cell suspension was centrifuged at 500×g for 5 minutes at 10 °C, and the supernatant was carefully removed without disturbing the cell pellet. The cells were resuspended in 100 μl red blood cell (RBC) lysis buffer (Cat# 11814389001, Roche) for the experiments involving two photobleaching fluorescent barcodes and incubated for 5 minutes at RT. The suspension was then centrifuged at 500×g for 5 minutes at 10 °C, and the supernatant was removed. The RBC lysis step was omitted in experiments involving a single fluorescence barcode on the Alexa Fluor 647 signal, as the autofluorescence impact from blood tends to be lower at this channel. Cell pellet was resuspended in 100 μl PBS, followed by adding 1 μl DAPI (5 mg/ml) to stain cell nuclei for 15 minutes at RT. Subsequently, 400 μl PBS was added, and the cell suspension was centrifuged at 500×g for 5 minutes at 10 °C. The supernatant was removed, and the final cell pellet was resuspended in 1% PFA and stored at 4 °C for FACS the following day.

### Fluorescence-activated cell sorting (FACS)

Fluorescent barcoded cells were sorted using the MACSQuant Tyto sorter with standard (4 ml/h flow rate) or high-speed (8 ml/h flow rate) cartridges. The temperature was maintained at 4 °C during cell sorting. Mouse splenocytes were diluted with PBS to a concentration of 1 million cells/ml and filtered through a 20 μm mesh (Cat# 07-001-105, Fisher Scientific). For every 10 ml of diluted cell solution, 40 μl of 2.5 mg/ml DNase (Cat# D5025, Sigma-Aldrich) was added and loaded into the sorting cartridge. For initial experiments, a “baseline” control (staining with DAPI only) and a ‘non-photobleaching’ control sample (staining with DAPI and fluorescent antibodies but without photobleaching) were prepared to establish the gating strategy. Unused channels were set to 300 V or lower to minimize background interference. After sorting, the target cell population was collected from the cartridge and transferred to a low-binding tube (Cat# 022431081, Eppendorf). The sample was kept on ice while sorting other groups. Lastly, the sorted cell solution was either processed immediately for protein extraction or stored at -80 °C for later analysis.

### Protein extraction

To lyse and extract proteins from the sorted cells, we prepared 10× RIPA lysis buffer (Cat# 20188, Sigma-Aldrich), 7× cOmplete protease inhibitor cocktail (dissolved in ddH_2_O) (Cat# 4693159001, Sigma-Aldrich), and 20× PhosSTOP phosphatase inhibitor cocktail (dissolved in ddH_2_O) (Cat# 4906845001, Sigma-Aldrich). These reagents were added directly to the sorted cell solution with volumes adjusted to achieve a final working concentration of 1× for each reagent. To enhance protein lysis, sodium dodecyl sulfate (SDS) (Cat# L3771, Sigma-Aldrich) was added to achieve a final concentration of 3% (w/v). The cell solution was incubated on a dry block at 95 °C for 30 minutes and then cooled to RT. Subsequently, the solution was further lysed at 65 °C for 1 hour with frequent vortexing, then cooled to RT again. DNase I and its 10× reaction buffer (Cat# 89836, Thermo Scientific) were added to the lysate at a 1:10 dilution ratio to the total volume of the sorted cell solution plus lysis buffer. The mixture was incubated at 37 °C for 30 minutes with frequent vortexing to degrade DNA and improve protein solubilization. Lastly, protein concentration and yield were quantified using a bicinchoninic acid (BCA) assay (Cat# 23227, Thermo Scientific).

### Protein digestion and liquid chromatography-mass spectrometry (LC-MS/MS)

Extracted and solubilized proteins from the sorted cell solution were mixed with an equal volume of 2× lysis buffer to a final concentration of 5% SDS (Cat# L3771, Sigma-Aldrich) and 50 mM triethanolamine bicarbonate (TEAB) (Cat#18597, Sigma-Aldrich). Tris(2-carboxyethyl)phosphine (TCEP) (Cat# 646547, Sigma-Aldrich) was added to the solution and incubated at 95 °C for 10 minutes to reduce disulfide bonds. Afterwards, iodoacetamide (IAA) (Cat# RPN6302, Cytiva) was added to the solution at a final concentration of 10 mM and incubated for 20 minutes under light protection to alkylate the free cysteines. The solution was then acidified with 2.5% phosphoric acid (Cat# 49685, Sigma-Aldrich), vortexed, and combined with binding/wash buffer (100 mM TEAB/90% MeOH). The sample was loaded onto an S-Trap column in 100 μl aliquots, ensuring the solution did not exceed the neck of the column. Each aliquot was centrifuged at 4,000×g for 30 seconds, and the process was repeated until the entire sample passed through the column. The S-Trap was washed three times with 200 µl of the binding/washing buffer, with each wash followed by centrifugation at 4,000×g for 30 seconds. After the final wash, the column was centrifuged for 1 minute at 4,000×g to ensure complete removal of residual liquid. Proteins retained on the S-trap were then digested overnight at 37 °C with trypsin (Cat# PRV5073, Fisher Scientific, Promega) at a protein-to-enzyme ratio of 10:1 (w/w). The next day, digested peptides were sequentially eluted into a clean low-binding tube using the following solutions, each centrifuged at 4,000×g for 1 minute: (1) 50 mM TEAB, pH 8.5; (2) 0.2% formic acid; (3) 50% acetonitrile/0.1% trifluoroacetic acid (TFA). The combined eluate was dried at -20 °C using a lyophilizer. Dried peptides were submitted to the Proteomics Core Facility at Mayo Clinic (Rochester, MN) for LC-MS/MS analysis.

Peptides were reconstituted in 40 µl sample buffer of 0.2% HCOOH, 0.1% TFA, 0.0005% z3-16, and 1 nM Pierce RST. After vortexing, the sample was spun for 1 minute at 10,000×g and transferred to autosampler vials. Peptide concentration was determined using Pierce Quantitative Fluorescence Peptide Assay (Cat# 23290, Thermo Scientific), and sample volumes were adjusted to ensure 140 ng of peptides were loaded across samples. LC-MS/MS was conducted on a Vanquish Neo UHPLC equipped with a trap column (Pep Map C18, 300 µm × 5 mm) and an IonOpticks Ultimate C18 column (1.7 µm; 75 µm × 25 cm) coupled to an Orbitrap Exploris 480 mass spectrometer (Thermo Scientific). Peptides were separated using a non-linear 145-minute gradient injecting at with 3% solvent B, increased to 35% over 120 minutes, ramped to 90% in 10 minutes, and held at 90% for an additional 10 minutes. Re-equilibration was performed at 3% solvent B for a duration three times the length of the column to restore starting conditions. Solvent A consisted of 0.1% formic acid and 2.5% acetonitrile in ddH_2_O, and solvent B was 0.1% formic acid, 80% acetonitrile, 10% isopropanol, and 10% water. The flow rate was maintained at 300 nL/min, and the column temperature was set to 50 °C.

The mass spectrometry data was acquired using a data dependent acquisition method with MS1 spectra acquired at 120,000 resolution over an m/z range of 350-1500, with the automatic gain control (AGC) target set at 200% and a maximum injection time of 50 ms. Precursors with charge states of 2+ to 6+ were isolated with a 1.2 Th isolation width and fragmented by higher-energy collisional dissociation (HCD) using a normalized collision energy (NCE) of 30. MS/MS spectra were collected at a resolution of 15,000 with an AGC target of 100%, a minimum intensity threshold of 5 × 10⁴, and a maximum injection time of 120 ms.

### Evaluation of protein artifacts in tissue cryopreservation

Mouse spleens were harvested and fixed in 4% PFA solution for 20 minutes at RT. The spleens were washed in cold PBS three times for 5 minutes each at 4 °C. Each spleen was bisected, with one half designated as the cryopreservation group and the other as the control. Cryopreserved spleens were incubated in 100% (w/v) D-fructose solution prepared in 100 mM PB buffer (pH 7.8) under gentle agitation for 1 hour at RT and stored at -80 °C for a week, while control spleens were dissociated into single-cell suspensions on the same day of harvesting. After one week, the cryopreserved spleens were thawed and washed in cold PBS four times for 5 minutes each at 4 °C to remove residual D-fructose. These tissues were then dissociated into single-cell suspensions following the same protocol described above. All resulting cell suspensions underwent protein extraction, peptide digestion, and LC-MS/MS analysis.

### Evaluation of protein artifacts in the NicheProt workflow

Mouse spleens were harvested, fixed in 4% PFA solution, and cryopreserved in 100% (w/v) D-fructose at -80 °C. After one week, the cryopreserved spleens were thawed at 4 °C and washed in cold PBS four times for 5 minutes each at 4 °C. Each spleen was bisected, with one half designated as the *NicheProt*-processed group and the other as the control group. Control spleens were sliced into 400 μm-thick macrosections and dissociated into single-cell suspensions on the same day. In contrast, spleens in the processed group underwent the *NicheProt* workflow, excluding the FACS step. In brief, the spleens were macrosectioned at 400 μm thickness and stained with DAPI (5 mg/ml, 1:500 v/v), Alexa Fluor 488-anti-CD45 (0.5 mg/ml, 1:20 v/v), and Alexa Fluor 647-anti-CD45 (0.5 mg/ml, 1:20 v/v) antibodies in 500 µl staining buffer composed of RPMI with 10 mg/ml BSA for 20 hours at 4 °C. The following day, spleen macrosections were washed, cleared with 50% and 80% (w/v) D-fructose solutions as described above, and photobleached in arbitrary regions by using a 20× air objective and a 640 nm excitation laser. The spleen macrosections were then reverse-cleared from D-fructose solution back into PBS and dissociated into single-cell suspensions following the same protocol as described above. All resulting cell suspensions underwent protein extraction, peptide digestion, and LC-MS/MS analysis.

### NicheProt analysis of DCs in the normal and inflamed mouse spleens

Spleens were harvested from healthy and LPS-treated mice and fixed in 4% PFA solution for 20 minutes at RT. The spleens were then washed three times in cold PBS for 5 minutes each at 4 °C and cryopreserved in 100% (w/v) D-fructose solution at -80 °C until analysis. On the day of the experiment, spleens were thawed and washed three times in cold PBS for 5 minutes each at 4 °C. The tissues were embedded in 2% agarose gel (LE Quick Dissolve Agarose, GeneMate) dissolved in distilled water and sliced into 400 µm-thick macrosections under cold conditions as described above. The macrosections were stained with 1 µl DAPI (5 mg/ml), 25 µl Alexa Fluor 488-anti-CD11c (0.5 mg/ml), 15 µl DyLight550-anti-CD3 (0.5 mg/ml), and 25 µl Alexa Fluor 647-anti-CD11c (0.5 mg/ml) antibodies in 500 µl staining buffer composed of RPMI with 10 mg/ml BSA. IF staining was carried out for 20 hours at 4 °C with gentle agitation. After staining, the macrosections were washed with PBS and cleared with 50% and 80% (w/v) D-fructose solutions to prepare for microscopic photobleaching. Fluorescence barcodes were generated in ROI regions within spleen macrosections by selectively photobleaching the Alexa Fluor 647 signal using a 20× air objective and a 640 nm excitation laser (intensity: 95%, duration: 20-40 seconds) on a RS-G4 confocal microscope (Caliber I.D.). Once barcoding was completed, the macrosections were reverse-cleared from D-fructose solution back into PBS and dissociated into the single-cell suspensions. FACS was performed to collect barcoded DC populations, followed by protein extraction, peptide digestion, and LC-MS/MS analysis.

### Image data analysis

Confocal microscopy images taken before and after photobleaching were processed and visualized using open-source Fiji software (http://fiji.sc/Fiji) and Imaris 10.1 software (Oxford Instrument, https://imaris.oxinst.com/). In Fiji, z-stack images of each channel were processed for brightness and contrast, and a Gaussian blur filter (radius=1) was applied to each channel before merging to a multicolor composite image. TIFF files were converted to .ims format using the Imaris File Converter (https://imaris.oxinst.com/microscopy-imaging-software-free-trial#file-converter) and imported into Imaris for further 3D rendering and visualization.

To estimate the number of splenocytes within photobleaching regions of varying volumes (**Table S1**), z-stack images were reconstructed in 3D using Fiji software, and DAPI^+^ nuclei were segmented with the Imaris “Surface” function. In Imaris, the photobleaching volume of each barcode was measured directly from the images, while the corresponding cell numbers were obtained from the segmentation result. Using the resulting cell number–to–volume ratios, we estimated the total number of cells across the 10 photobleaching regions generated with different objectives and zoom factors

### Proteomics data analysis

Mass spectrometer (MS) raw files were searched against a Swissprot mouse database (SP_mouse_2024_06; 17208 entries) using the Sequest HT node in Proteome Discoverer Software (version 3.0.0.757). The search parameters were set for full trypsin specificity, allowing 3 missed cleavages with oxidized Met and N-term protein acetylation as variable modifications and carbamidomethyl cysteine as fixed modification. Mass tolerances were set at 10 ppm for precursor ions and 0.02 Dalton for MS2 fragment ions. High confidence peptide spectral matches (PSMs) are assigned using the Percolator node and peptide and protein identifications are filtered at a 1% false discovery rate (FDR). Intensity-based absolute quantification (iBAQ) values for each protein were calculated from summed ion peak abundances of the matched peptides divided by the number of tryptic peptides theoretically detectable with the mass spectrometry acquisition parameters. The iBAQ values represent a molar abundance estimate that was used for comparing protein amounts between samples. A blank iBAQ value indicates that protein abundance was below quantitation levels or matched peptides were shared with another protein.

Proteomics downstream data analysis and visualization were performed using R (version 2024.09.1+394). The filtering criterion was applied to retain only the quantified protein groups with at least two valid values in at least one group. Data normalization was done using a variance-stabilizing normalization (VSN) package. A MeanSD plot was generated in R using the ggplot2 package to evaluate the data distribution and ensure the quality control of the datasets. Missing values were imputed from a normal distribution with a width of 0.3 standard deviations, downshifted by 1.8 standard deviations. Batch effect correction was applied in processing datasets of “Protein artifact evaluation in D-fructose cryopreservation” and “*NicheProt* analysis of DCs in the normal and inflamed mouse spleens” using the ComBat method. Proteome correlations across samples or biological replicates were performed after imputation, normalization, and batch correction. Coefficients of variation (CVs) were calculated within experimental groups using imputed and normalized datasets. A principal component analysis (PCA) plot was computed using the method “singular value decomposition (svd)” and visualized in ggplot2. A variance protein heatmap was generated based on the top 50 proteins with significant variation across different groups with normalized rows to better visualize relative expression (scale: “row”). A hierarchical clustering heatmap was computed using the Euclidean method to calculate the differences across samples and compare the clusters with the complete-linkage method. Differential gene expression analysis was performed using the limma and edgeR packages and visualized in ggplot2. A linear model is fitted for each gene, and the *p*-value is computed with a moderated t-test with empirical Bayes moderation in limma for differential analysis. Benjamini-Hochberg correction was applied to control the false discovery rate for multiple testing corrections.

## Acknowledgements

We thank the Mayo Clinic Proteomics Core for providing mass spectrometry services and valuable discussions. We also thank the Proteomics Core at the University of Chicago for their consultation and recommendations on sample preparation. We are grateful to Dr. Ryan Wilcox and Dr. Nermin Kady at University of Michigan for providing tissue samples used in the pilot testing of our technology. We acknowledge the use of ChatGPT-4.0 for text editing suggestions and assistance in revising the manuscript written by the authors.

This work was supported by NIGMS R35 GM142743 (SSYL), NCI R21 CA286296 (SSYL), the University of Illinois Cancer Center Pilot Project Awards 2020-PP-07 and 2023-29-UICCPG (SSYL), NHLBI R01 HL174916 (JR) and P01 HL160469 (JR), and NCI R33 CA258012 (JR).

## Author contribution

Y.-C.W. and S.S.-Y.L. conceived and designed the study. Y.-C.W. conducted the experiments, performed data analysis, and wrote the manuscript. D.S. and E.A.K. contributed to sample preparation and manuscript revision. A.U. developed the data analysis pipeline and performed data analysis. J.R. contributed to project conceptualization and data interpretation. S.S.-Y.L. supervised the study and revised the manuscript. All authors provided input, edited, and approved the final version of the manuscript.

## Supplementary Figure Legends

**Figure S1.**
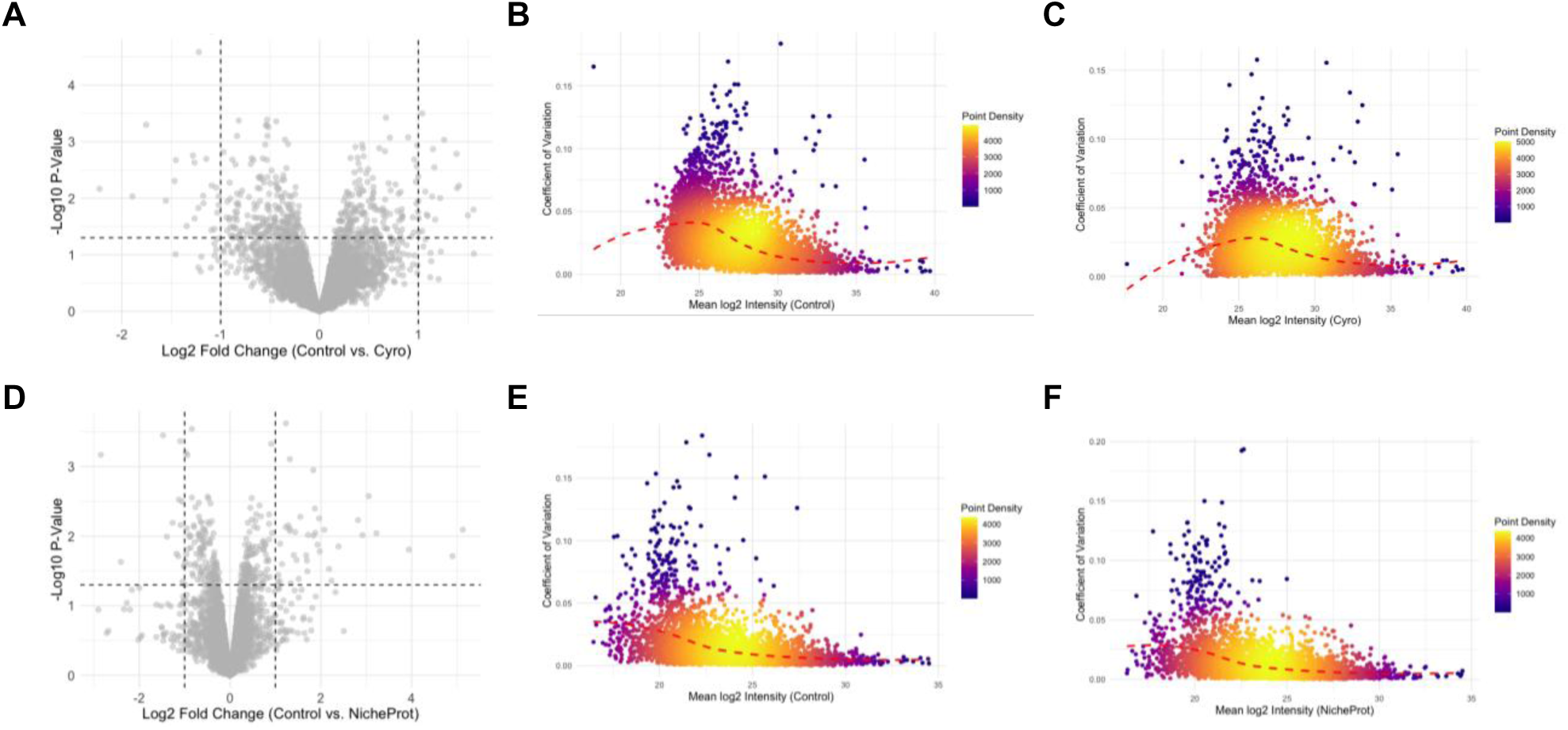
Quantitative assessment of the proteome in control and NicheProt samples. (A) Volcano plot comparing control and cryopreserved samples. (B, C) Coefficients of variation (CVs) within control (B) and cryopreservation (C) groups were below 0.2, demonstrating high reproducibility between biological replicates. (D) Volcano plot comparing control and NicheProt samples showed no significant protein expression changes. (E, F) CVs plot within the control (E) and NicheProt (F) conditions were below 0.2.

**Figure S2.**
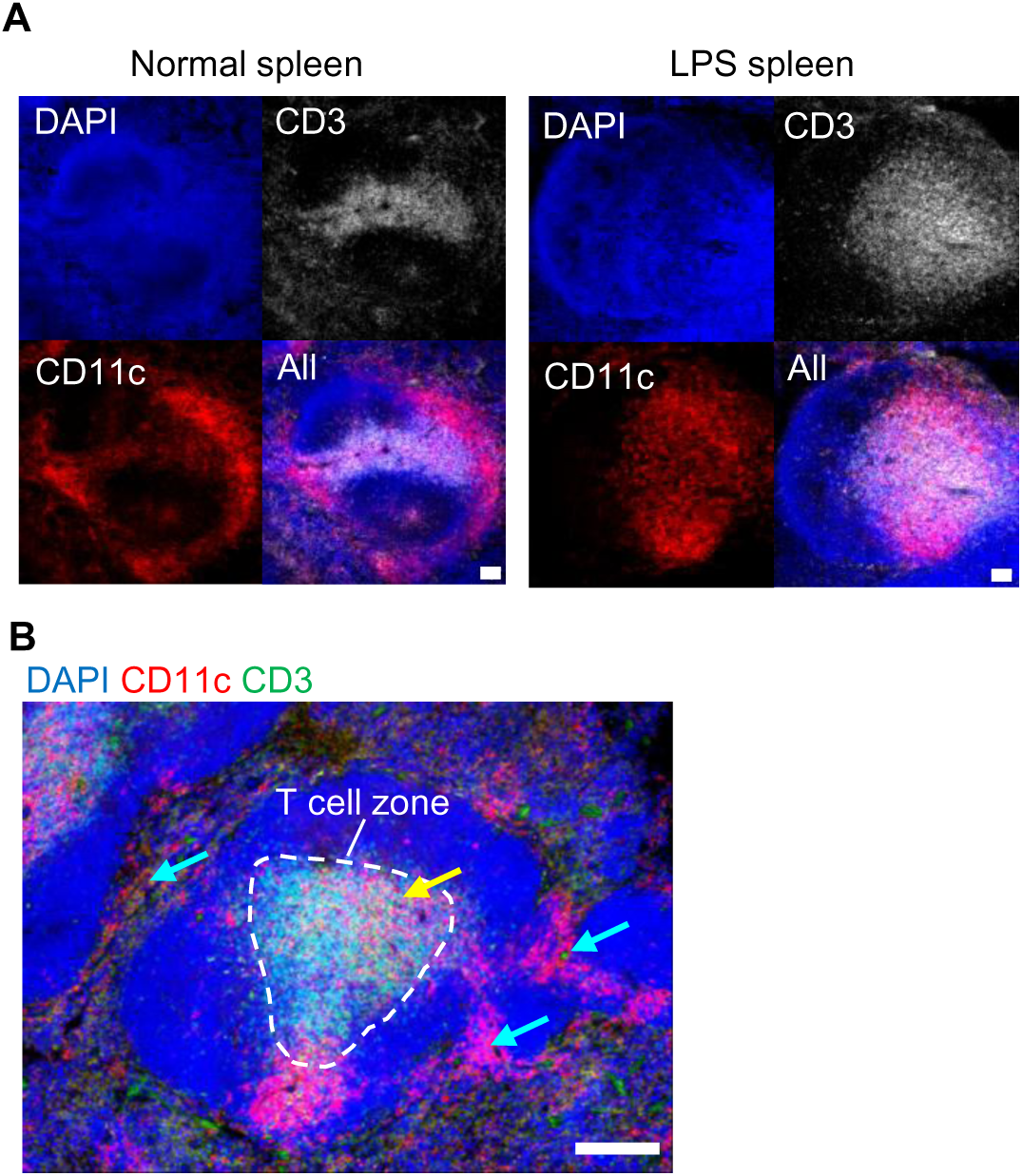
Dendritic cell migration during LPS-induced inflammation. (A) Splenic dendritic cells (DCs) primarily reside outside the T cell zone (TCZ) in a normal spleen. DCs migrate toward the TCZ upon LPS-induced inflammation. DCs were labeled with Alexa Fluor 647-anti-CD11c antibody, while T cells were stained with DyLight 550-anti-CD3 antibody. Scale bars: 50 µm. (B) During inflammation, there are two DC populations located *inside* (yellow arrow) and *outside* (cyan arrows) the TCZ. Scale bar: 100 µm.

**Figure S3.**
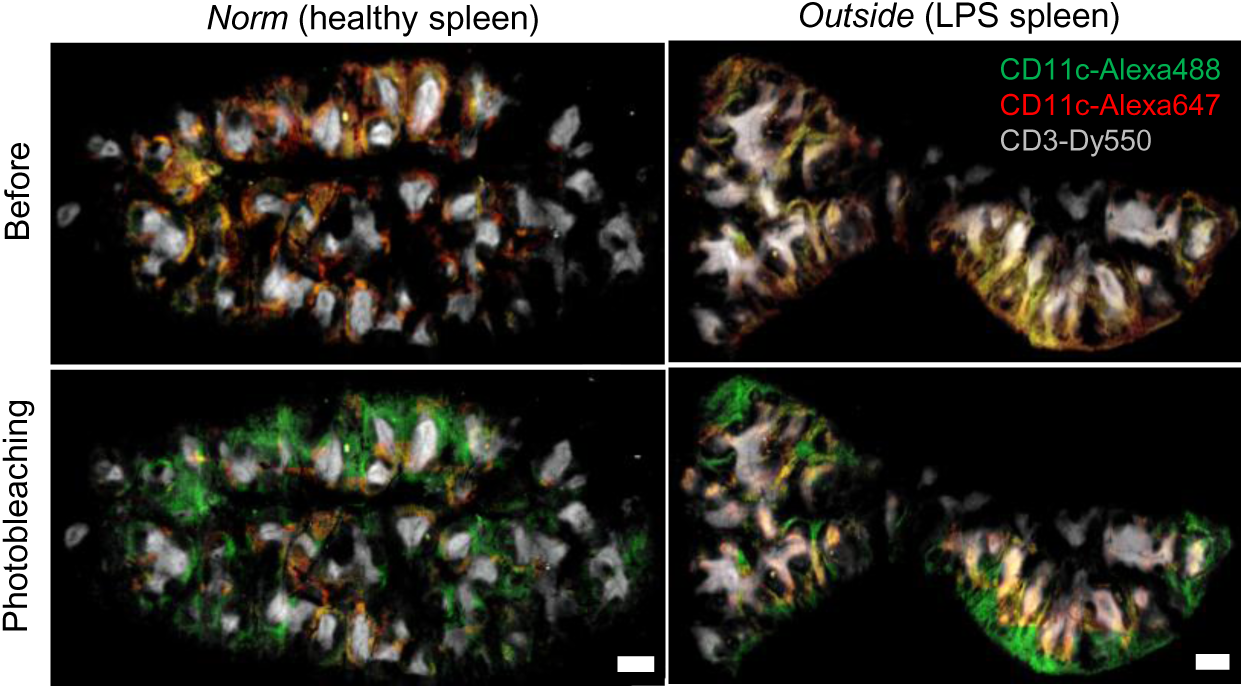
Photobleaching-mediated barcoding of DCs outside the TCZs in the (left) healthy and (right) LPS-treated mouse spleen macrosections. Tissue macrosections were labeled with Alexa Fluor 488-anti-CD11c (green), Alexa Fluor 647-anti-CD11c (red), and Dylight-CD3 (gray). Photobleaching of Alexa Fluor 647 on CD11c⁺ DCs outside the CD3⁺ TCZs converted yellow signals to green. Scale bar: 500 µm.

**Table S1.** The volumes, estimated cell numbers, and protein amounts of 3D photobleached regions in mouse spleen macrosections using 20× and 40× objectives with varying zoom factors.

| Objective | Zoom factor | x (μm) | y (μm) | z (μm) | Volume (μm³) | Estimated cell number (10 areas) | Estimated protein (μg) (10 areas) |
| --- | --- | --- | --- | --- | --- | --- | --- |
| 20x air | 1.5x | 610 | 500 | 400 | 1.22E+08 | 4.12E+05 | 12.31 |
|  | 2x | 450 | 400 | 400 | 7.20E+07 | 2.43E+05 | 7.26 |
|  | 3x | 300 | 260 | 400 | 3.12E+07 | 1.05E+05 | 3.14 |
|  | 4x | 220 | 225 | 400 | 1.98E+07 | 6.69E+04 | 2.00 |
| 40x air | 3x | 230 | 170 | 400 | 1.56E+07 | 5.29E+04 | 1.58 |
|  | 4x | 125 | 110 | 400 | 5.50E+06 | 1.86E+04 | 0.56 |

**Table S2.**
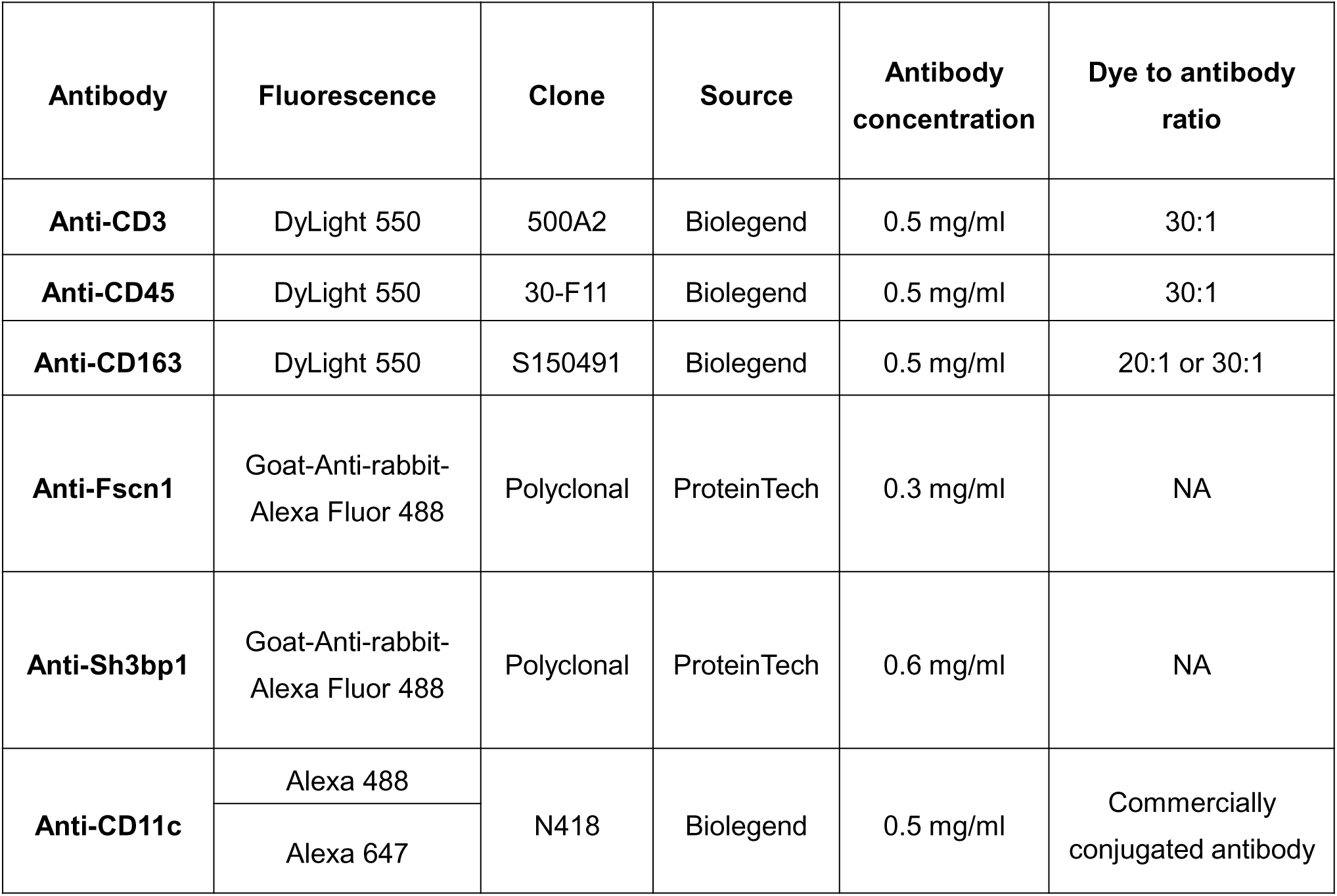
Antibody information.

## Notes

### Competing Interest Statement

The authors have declared no competing interest.

## References

1. Palla, G., et al., Spatial components of molecular tissue biology. Nat Biotechnol, 2022. 40(3): p. 308–318.

2. Lázár, E. and J. Lundeberg, Spatial architecture of development and disease. Nat Rev Genet, 2025. 10.1038/s41576-025-00892-5

3. Liu, L., et al., Spatiotemporal omics for biology and medicine. Cell, 2024. 187(17): p. 4488–4519.

4. Park, J., et al., Spatial omics technologies at multimodal and single cell/subcellular level. Genome Biol, 2022. 23(1): p. 256.

5. You, Y., et al., Systematic comparison of sequencing-based spatial transcriptomic methods. Nat Methods, 2024. 21(9): p. 1743–1754.

6. Coleman, K., et al., Resolving tissue complexity by multimodal spatial omics modeling with MISO. Nat Methods, 2025. 22(3): p. 530–538.

7. Pei, G., et al., Spatial mapping of transcriptomic plasticity in metastatic pancreatic cancer. Nature, 2025. 642(8066): p. 212–221.

8. Liu, Y., et al., Conserved spatial subtypes and cellular neighborhoods of cancer-associated fibroblasts revealed by single-cell spatial multi-omics. Cancer Cell, 2025. 43(5): p. 905–924.e6.

9. Gaspard-Boulinc, L.C., et al., Cell-type deconvolution methods for spatial transcriptomics. Nat Rev Genet, 2025. 10.1038/s41576-025-00845-y

10. Takemon, Y., et al., Proteomic and transcriptomic profiling reveal different aspects of aging in the kidney. Elife, 2021. 10: p. e62585.

11. Chen, G., et al., Discordant protein and mRNA expression in lung adenocarcinomas. Mol Cell Proteomics, 2002. 1(4): p. 304–13.

12. Landgraf, A., et al., Widespread discordance between mRNA expression, protein abundance and de novo lipogenesis activity in hepatocytes during the fed-starvation transition. bioRxiv, 2025. doi: 10.1101/2025.04.15.649020.

13. Keren, L., et al., MIBI-TOF: A multiplexed imaging platform relates cellular phenotypes and tissue structure. Sci Adv, 2019. 5(10): doi: 10.1126/sciadv.aax585

14. Angelo, M., et al., Multiplexed ion beam imaging of human breast tumors. Nat Med, 2014. 20(4): p. 436–42.

15. Angelo, M., et al., Quantitative Imaging of Ten Markers in Human Breast Tumors Using Multiplexed Ion Beam Imaging (MIBI) and Metal-Tagged Antibodies. Modern Pathol, 2014. 27(518): p. 436–442.

16. Giesen, C., et al., Highly multiplexed imaging of tumor tissues with subcellular resolution by mass cytometry. Nat Methods, 2014. 11(4): p. 417–22.

17. Black, S., et al., CODEX multiplexed tissue imaging with DNA-conjugated antibodies. Nat Protoc, 2021. 16(8): p. 3802–3835.

18. Kuswanto, W., G. Nolan, and G. Lu, Highly multiplexed spatial profiling with CODEX: bioinformatic analysis and application in human disease. Semin Immunopathol, 2023. 45(1): p. 145–157.

19. Liu, Y., et al., High-Spatial-Resolution Multi-Omics Sequencing via Deterministic Barcoding in Tissue. Cell, 2020. 186(6): p. 1665–1681.

20. Merritt, C.R., et al., Multiplex digital spatial profiling of proteins and RNA in fixed tissue. Nat Biotechnol, 2020. 38(5): p. 586–599.

21. Unterauer, E.M., et al., Spatial proteomics in neurons at single-protein resolution. Cell, 2024. 187(7): p. 1785–1800.e16.

22. Zheng, J., et al., Increased Multiplexity in Optical Tissue Clearing-Based Three-Dimensional Immunofluorescence Microscopy of the Tumor Microenvironment by Light-Emitting Diode Photobleaching. Lab Invest, 2024. 104(6): p. 102072.

23. Kinkhabwala, A., et al., MACSima imaging cyclic staining (MICS) technology reveals combinatorial target pairs for CAR T cell treatment of solid tumors. Sci Rep, 2022. 12(1): p. 1911.

24. Lin, J.R., et al., Highly multiplexed immunofluorescence imaging of human tissues and tumors using t-CyCIF and conventional optical microscopes. Elife, 2018. 7: p. e31657.

25. Makhmut, A., et al., A framework for ultra-low-input spatial tissue proteomics. Cell Syst, 2023. 14(11): p. 1002–1014.e5.

26. Mund, A., A.D. Brunner, and M. Mann, Unbiased spatial proteomics with single-cell resolution in tissues. Mol Cell, 2022. 82(12): p. 2335–2349.

27. Mongia, A., et al., AnnoSpat annotates cell types and quantifies cellular arrangements from spatial proteomics. Nat Commun, 2024. 15(1): p. 3744.

28. Wu, Y.C., et al., Multiresolution 3D Optical Mapping of Immune Cell Infiltrates in Mouse Asthmatic Lung. Am J Respir Cell Mol Biol, 2023. 69(1): p. 13–21.

29. Zheng, J., et al., Correlative multiscale 3D imaging of mouse primary and metastatic tumors by sequential light sheet and confocal fluorescence microscopy. iScience, 2025. 28(3): p. 111934.

30. Choudhury, F.K., et al., Multiomics Characterization of a Less Invasive Microfluidic-Based Cell Sorting Technique. J Proteome Res, 2024. 23(8): p. 3096–3107.

31. Staunstrup, N.H., et al., Comparison of electrostatic and mechanical cell sorting with limited starting material. Cytometry A, 2022. 101(4): p. 298–310.

32. Lewis, S.M., A. Williams, and S.C. Eisenbarth, Structure and function of the immune system in the spleen. Sci Immunol, 2019. 4(33). doi: 10.1126/sciimmunol.aau6085.

33. Bronte, V. and M.J. Pittet, The spleen in local and systemic regulation of immunity. Immunity, 2013. 39(5): p. 806–18.

34. Backer, R.A., N. Diener, and B.E. Clausen, Langerin ^+^ CD8 ^+^ Dendritic Cells in the Splenic Marginal Zone: Not So Marginal After All. Front Immunol, 2019. 10: p. 741.

35. Goltsev, Y., et al., Deep Profiling of Mouse Splenic Architecture with CODEX Multiplexed Imaging. Cell, 2018. 174(4): p. 968–981.e15.

36. Villadangos, J.A., P. Schnorrer, and N.S. Wilson, Control of MHC class II antigen presentation in dendritic cells: a balance between creative and destructive forces. Immunol Rev, 2005. 207: p. 191–205.

37. Benvenuti, F., et al., Requirement of Rac1 and Rac2 expression by mature dendritic cells for T cell priming. Science, 2004. 305(5687): p. 1150–3.

38. Calabro, S., et al., Differential Intrasplenic Migration of Dendritic Cell Subsets Tailors Adaptive Immunity. Cell Rep, 2016. 16(9): p. 2472–85.

39. Alvarez, D., E.H. Vollmann, and U.H. von Andrian, Mechanisms and consequences of dendritic cell migration. Immunity, 2008. 29(3): p. 325–42.

40. Xiao, Q. and Y. Xia, Insights into dendritic cell maturation during infection with application of advanced imaging techniques. Front Cell Infect Microbiol, 2023. 13: p. 1140765.

41. Feng, M., et al., Regulation of the Migration of Distinct Dendritic Cell Subsets. Front Cell Dev Biol, 2021. 9: p. 635221.

42. Xu, L., et al., Time-dependent effect of E. coli LPS in spleen DC activation in vivo: Alteration of numbers, expression of co-stimulatory molecules, production of pro-inflammatory cytokines, and presentation of antigens. Mol Immunol, 2017. 85: p. 205–213.

43. Seemann, S., F. Zohles, and A. Lupp, Comprehensive comparison of three different animal models for systemic inflammation. J Biomed Sci, 2017. 24(1): p. 60.

44. Dervishi, E., et al., Early-Life Exposure to Lipopolysaccharide Induces Persistent Changes in Gene Expression Profiles in the Liver and Spleen of Female FVB/N Mice. Vet Sci, 2023. 10(7): p. 445.

45. Jiao, Y., et al., Intraperitoneal versus intranasal administration of lipopolysaccharide in causing sepsis severity in a murine model: a preliminary comparison. Lab Anim Res, 2024. 40(1): p. 18.

46. Dixon, G.L., et al., Dendritic cell activation and cytokine production induced by group B Neisseria meningitidis: interleukin-12 production depends on lipopolysaccharide expression in intact bacteria. Infect Immun, 2001. 69(7): p. 4351–7.

47. Lee, K.W., et al., Direct role of NF-kappaB activation in Toll-like receptor-triggered HLA-DRA expression. Eur J Immunol, 2006. 36(5): p. 1254–66.

48. Arya, S., et al., Quantitative proteomic changes in LPS-activated monocyte-derived dendritic cells: A SWATH-MS study. Sci Rep, 2019. 9(1): p. 4343.

49. Singh, P. and S.A. Ali, Multifunctional Role of S100 Protein Family in the Immune System: An Update. Cells, 2022. 11(15): p. 2274.

50. Sreejit, G., et al., S100 family proteins in inflammation and beyond. Adv Clin Chem, 2020. 98: p. 173–231.

51. Lim, R.J., et al., CXCL9/10-engineered dendritic cells promote T cell activation and enhance immune checkpoint blockade for lung cancer. Cell Rep Med, 2024. 5(4): p. 101479.

52. Tokunaga, R., et al., CXCL9, CXCL10, CXCL11/CXCR3 axis for immune activation - A target for novel cancer therapy. Cancer Treat Rev, 2018. 63: p. 40–47.

53. Marquet, J., et al., Dichotomy between factors inducing the immunosuppressive enzyme IL-4-induced gene 1 (IL4I1) in B lymphocytes and mononuclear phagocytes. Eur J Immunol, 2010. 40(9): p. 2557–68.

54. Chatterjee, P., et al., Regulation of the Anti-Inflammatory Cytokines Interleukin-4 and Interleukin-10 during Pregnancy. Front Immunol, 2014. 5: p. 253.

55. Puiffe, M.L., et al., IL4I1 Accelerates the Expansion of Effector CD8. Front Immunol, 2020. 11: p. 600012.

56. Shimizu, K., et al., Granzyme A Stimulates pDCs to Promote Adaptive Immunity via Induction of Type I IFN. Front Immunol, 2019. 10: p. 1450.

57. van Eck, J.A., et al., A novel proinflammatory role for granzyme A. Cell Death Dis, 2017. 8(2): p. e2630.

58. Velotti, F., et al., Granzyme B in Inflammatory Diseases: Apoptosis, Inflammation, Extracellular Matrix Remodeling, Epithelial-to-Mesenchymal Transition and Fibrosis. Front Immunol, 2020. 11: p. 587581.

59. Wu, R., et al., The Dual Role of ACOD1 in Inflammation. J Immunol, 2023. 211(4): p. 518–526.

60. Wu, R., et al., ACOD1 in immunometabolism and disease. Cell Mol Immunol, 2020. 17(8): p. 822–833.

61. Wu, R., R. Kang, and D. Tang, Mitochondrial ACOD1/IRG1 in infection and sterile inflammation. J Intensive Med, 2022. 2(2): p. 78–88.

62. Aziz, M.H., et al., The Upregulation of Integrin α D β 2 (CD11d/CD18) on Inflammatory Macrophages Promotes Macrophage Retention in Vascular Lesions and Development of Atherosclerosis. J Immunol, 2017. 198(12): p. 4855–4867.

63. Bailey, W.P., et al., The expression of integrin α D β 2 (CD11d/CD18) on neutrophils orchestrates the defense mechanism against endotoxemia and sepsis. J Leukoc Biol, 2021. 109(5): p. 877–890.

64. Zimmer, A., et al., A regulatory dendritic cell signature correlates with the clinical efficacy of allergen-specific sublingual immunotherapy. J Allergy Clin Immunol, 2012. 129(4): p. 1020–30.

65. Shen, M., et al., Tinagl1 Suppresses Triple-Negative Breast Cancer Progression and Metastasis by Simultaneously Inhibiting Integrin/FAK and EGFR Signaling. Cancer Cell, 2019. 35(1): p. 64–80.e7.

66. Grin, P.M., et al., Low-density lipoprotein (LDL)-dependent uptake of Gram-positive lipoteichoic acid and Gram-negative lipopolysaccharide occurs through LDL receptor. Sci Rep, 2018. 8(1): p. 10496.

67. Radford-Smith, D.E., et al., HDL and LDL have distinct, opposing effects on LPS-induced brain inflammation. Lipids Health Dis, 2023. 22(1): p. 54.

68. Arnob, A., et al., Factors Promoting Lipopolysaccharide Uptake by Synthetic Lipid Droplets. ACS Omega, 2025. 10(6): p 5866–5873.

69. Sen Chaudhuri, A., et al., S100A4 exerts robust mucosal adjuvant activity for co-administered antigens in mice. Mucosal Immunol, 2022. 15(5): p. 1028–1039.

70. Li, Z.H., et al., S100A4 regulates macrophage chemotaxis. Mol Biol Cell, 2010. 21(15): p. 2598–610.

71. Siegfried, A., et al., IFIT2 is an effector protein of type I IFN-mediated amplification of lipopolysaccharide (LPS)-induced TNF-α secretion and LPS-induced endotoxin shock. J Immunol, 2013. 191(7): p. 3913–21.

72. Hong, L., T.J. Webb, and D.S. Wilkes, Dendritic cell-T cell interactions: CD8 alpha alpha expressed on dendritic cells regulates T cell proliferation. Immunol Lett, 2007. 108(2): p. 174–8.

73. Karigane, D., et al., Mitf is required for T cell maturation by regulating dendritic cell homing to the thymus. Biochem Biophys Res Commun, 2022. 596: p. 29–35.

74. Glatigny, S., et al., Cutting edge: loss of α4 integrin expression differentially affects the homing of Th1 and Th17 cells. J Immunol, 2011. 187(12): p. 6176–9.

75. Ryu, S., et al., The protective roles of integrin α4β7 and Amphiregulin-expressing innate lymphoid cells in lupus nephritis. Cell Mol Immunol, 2024. 21(7): p. 723–737.

76. Mylvaganam, S., S.A. Freeman, and S. Grinstein, The cytoskeleton in phagocytosis and macropinocytosis. Curr Biol, 2021. 31(10): p. R619–R632.

77. Frittoli, E., et al., The signaling adaptor Eps8 is an essential actin capping protein for dendritic cell migration. Immunity, 2011. 35(3): p. 388–99.

78. Elbediwy, A., et al., Epithelial junction formation requires confinement of Cdc42 activity by a novel SH3BP1 complex. J Cell Biol, 2012. 198(4): p. 677–93.

79. Medaglia, C., et al., Spatial reconstruction of immune niches by combining photoactivatable reporters and scRNA-seq. Science, 2017. 358(6370): p. 1622–1626.

80. Schaupp, L., et al., Microbiota-Induced Type I Interferons Instruct a Poised Basal State of Dendritic Cells. Cell, 2020. 181(5): p. 1080–1096.e19.

81. Jang, J.S., et al., Rsad2 is necessary for mouse dendritic cell maturation via the IRF7-mediated signaling pathway. Cell Death Dis, 2018. 9(8): p. 823.

82. Liu, X.Y., et al., IFN-induced TPR protein IFIT3 potentiates antiviral signaling by bridging MAVS and TBK1. J Immunol, 2011. 187(5): p. 2559–68.

83. Xu, F., et al., IFIT3 mediated the type I interferon antiviral response by targeting Senecavirus A entry, assembly and release pathways. Vet Microbiol, 2022. 275: p. 109594.

84. Harioudh, M.K., et al., The canonical antiviral protein oligoadenylate synthetase 1 elicits antibacterial functions by enhancing IRF1 translation. Immunity, 2024. 57(8): p. 1812–1827.e7.

85. Martin-Fernandez, M. and D. Bogunovic, A non-canonical function of OAS1 to combat viral and bacterial infections. Immunity, 2024. 57(8): p. 1721–1723.

86. Shenoy, A.R., et al., GBP5 promotes NLRP3 inflammasome assembly and immunity in mammals. Science, 2012. 336(6080): p. 481–5.

87. Marinho, F.V., et al., Guanylate-binding protein-5 is involved in inflammasome activation by bacterial DNA but only the cooperation of multiple GBPs accounts for control of. Front Immunol, 2024. 15: p. 1341464.

88. Yamakita, Y., et al., Fascin1 promotes cell migration of mature dendritic cells. J Immunol, 2011. 186(5): p. 2850–9.

89. Parrini, M.C., et al., SH3BP1, an exocyst-associated RhoGAP, inactivates Rac1 at the front to drive cell motility. Mol Cell, 2011. 42(5): p. 650–61.

90. Ballegeer, M., et al., Glucocorticoid receptor dimers control intestinal STAT1 and TNF-induced inflammation in mice. J Clin Invest, 2018. 128(8): p. 3265–3279.

91. Cotterell, J., et al., Cell 3D Positioning by Optical encoding (C3PO) and its application to spatial transcriptomics. bioRxiv, 2024. doi: 10.1101/2024.03.12.584578

92. Woo, J., et al., High-throughput and high-efficiency sample preparation for single-cell proteomics using a nested nanowell chip. Nat Commun, 2021. 12(1): p. 6246.

93. Matzinger, M., R.L. Mayer, and K. Mechtler, Label-free single cell proteomics utilizing ultrafast LC and MS instrumentation: A valuable complementary technique to multiplexing. Proteomics, 2023. 23(13-14): p. e2200162.

94. Brunner, A.D., et al., Ultra-high sensitivity mass spectrometry quantifies single-cell proteome changes upon perturbation. Mol Syst Biol, 2022. 18(3): p. e10798.

95. Walker, C., et al., PatchSorter: a high throughput deep learning digital pathology tool for object labeling. NPJ Digit Med, 2024. 7(1): p. 164.

96. Hossain, M.S., et al., Region of interest (ROI) selection using vision transformer for automatic analysis using whole slide images. Sci Rep, 2023. 13(1): p. 11314.

97. Xu, Y., et al., Multimodal single cell-resolved spatial proteomics reveal pancreatic tumor heterogeneity. Nat Commun, 2024. 15(1): p. 10100.

